# Exploring operational requirements of mixtures for insecticide resistance management in public health using a mathematical model assuming polygenic resistance

**DOI:** 10.1101/2024.05.06.592650

**Authors:** Neil Philip Hobbs, Ian Michael Hastings

## Abstract

Long-lasting insecticide treated nets (LLINs) have been developed which contain two active ingredients. Mixture products for vector control are now an available insecticide resistance management (IRM) strategy. There is a theoretical concern around the use mixtures pertaining to dosing, insecticide decay, initial resistance, and cross resistance. Mixture LLINs all currently have a pyrethroid as one of the partner insecticides. Using previously described mathematical models of polygenic insecticide resistance evolution, which implement selection either by truncation (“polytrucate) or as a probabilistic process (“polysmooth”) mixtures are evaluated for their IRM potential. Scenarios are developed to explore the impact of the initial levels of resistance to the pyrethroid insecticide, insecticide decay rates, and insecticide doses, and cross resistance. Results from our simulations indicate that mixtures should be deployed at full doses. As the initial level of resistance to the pyrethroid increases the benefit of mixture decreases. Insecticide decay was found to be less important than might be thought, with other variables having a greater impact. The mechanism of selection demonstrated consistent results, diverging only at very high levels of resistance to the pyrethroid. Our simulations demonstrate that the impact of positive cross resistance is best mitigated using full-dose mixtures. Insecticide decay less important than previously considered, and opens up the opportunity to mix a wider variety of insecticides. However, as mixtures remain a challenge to develop and therefore strategies which could generate an effect of a “temporal” mixture should be evaluated.

## 1. Introduction

Insecticides play a major role in the control of malaria (Bhatt et al., 2015). Until recently pyrethroids were the only insecticide available for use on long-lasting insecticide-treated nets (LLINs). This reliance on pyrethroids has led to widespread resistance (Hancock et al., 2020). The Global Plan for Insecticide Resistance Management (WHO, 2012) was designed to mitigate, slow or prevent, the spread of resistance and noted at the time of its development that mixtures while currently unavailable for vector control may become a potential future insecticide resistance management (IRM) strategy as alternatives to pyrethroids become available. Mixture products are now becoming available for use in vector control programmes and there is now an urgent need to evaluate their IRM capability within different deployment contexts. Next-generation mixture long-lasting insecticide treated nets (LLINs) are showing good epidemiological efficacy against malaria in field trials (Mosha et al., 2022; Tiono et al., 2018), however questions remain regarding their long term potential as IRM strategies. It is critical to evaluate the IRM implication of mixtures used in vector control especially in the context of their use, which at the current time is primarily as next-generation mixture LLINs which contain both a pyrethroid and a novel insecticide.

Mixtures are defined as a single formulation containing two insecticides, such that any mosquito encountering the formulation inevitably encounters both insecticides. Mixtures are considered a promising IRM strategy based on theoretical modelling studies performed over the last 40 years (e.g. Curtis, 1985; Madgwick & Kanitz, 2022; Mani, 1985; South & Hastings, 2018). However this conclusion requires certain conditions are met, notably: resistance to both insecticide partners is low; there is no cross resistance between the insecticides; the insecticides have similar residual lifespans; a portion of the target population remains unexposed to the mixture; and both insecticides in the mixture are at their respective full dose (Rex Consortium, 2013).

The requirement for resistance to each insecticide to be initially low is challenged by next-generation mixture LLINs using a pyrethroid as one of the partners (for example Interceptor G2® LLIN mixes chlorfenapyr and alpha-cypermethrin, whereas PermaNet Dual® LLIN mixes chlorfenapyr and deltamethrin). There is likely to be substantial resistance to the pyrethroid (Hancock et al., 2020), highlighting theoretical requirements not coinciding with practical realities.

Cross resistance occurs where some resistance to one insecticides impacts the resistance status to a another insecticide (Tabashnik et al., 2014). This can be positive cross resistance where the resistance for one insecticide also confers resistance to a different insecticide, or negative cross resistance where the resistance to one insecticide increases the susceptibility to another insecticide. Cross resistance is frequently absent from models evaluating IRM strategies (Rex Consortium, 2010) and therefore this issue is rarely evaluated, yet has been consistently highlighted as an issue (Curtis, 1985).

Insecticides deployed as LLINs or indoor residual sprays (IRS) decay over their deployed lifespan, which is especially important for vector control where the time between replacement or re-application can be 3 years for LLINs and 12 months for IRS. For mixtures, a concern often highlighted is the need for the two insecticides to have similar decay rates (Curtis, 1985). Insecticide decay has been noted to be frequently absent from models evaluating IRM strategies (South et al., 2020) and the importance of insecticide decay has been previously highlighted in simplistic scenarios (Hobbs & Hastings, 2024a).

One criterion for mixtures to remain effective is for a portion of the target population to remain unexposed to the mixture (Rex Consortium, 2013). This could be in the form of incomplete coverage in the target location or an influx of more susceptible individuals from untreated refugia (Comins, 1977). This means highly resistant individuals who survive the contact with the mixture are diluted by a larger number of more susceptible individuals. Of course, this likely comes with a trade-off between effective disease control and the time to insecticide failure (Sisterson, 2022).

A reduction in the dosage when insecticides are used in mixture is not an unreasonable expectation, and was an issue highlighted by Curtis (1985). Full doses of both insecticides in a mixture are likely expensive and/or complex to produce. Insecticide manufacturers may therefore reduce the dosage of one or both insecticides in the mixture to develop an economically, or operationally viable product. For some current mixture LLINs the dosage of the pyrethroid partner is lower than its standard-LLIN comparator. While new mixture products may be at lower doses than their single insecticide comparators, improved chemical technologies and manufacturing processes may mean reductions in dosages do not necessarily lead to a reduction in insecticide bioavailability and insecticidal ability. The IRM implications of reducing the dose in mixtures requires investigation (Rex Consortium, 2013).

Evaluating the efficacy or IRM strategies to delay resistance primarily rests on modelling as empirical evaluation is expensive, time-consuming, and cannot be realistically replicated over a wide range of epidemiological and operational settings. The purpose of this paper is therefore to evaluate how the deployment of mixtures in the form of next-generation LLINs perform as part of an IRM strategy across the range of highlighted required conditions. These highlighted conditions come primarily from models assuming resistance is monogenic (Rex Consortium, 2013) and the important question is do these requirement extend to polygenic resistance? Using previously described models (Hobbs & Hastings, 2024b), we explore these conditions and discuss the implication of these conditions not being met.

## 2. Methods

A previously detailed model of insecticide selection is used which allows for the modelling of polygenic resistance (Hobbs & Hastings, 2024b). Simulations are separated into two scenarios. Scenario 1 explores the impact of the level of pre-existing resistance, insecticide dosing and insecticide decay on the IRM capability of mixtures. Scenario 2 explores the impact of insecticide dosing and cross resistance on the IRM capability of mixtures.

### 2.1. Model Overview

A full detailed description of the mathematical model is given elsewhere (Hobbs & Hastings, 2024b), but key aspects are highlighted here: it is a quantitative genetics model which tracks a “polygenic resistance score” (PRS) for each insecticide which quantifies the “amount of resistance” and is a classically quantitative trait. The PRS is measurable in standardised bioassays (e.g., WHO cylinder). The selection differentials are the within generation change in the mean PRS and are calculated separately for males and females. The selection differentials are calculated dynamically each generation and are dependent on the current amount of resistance in the mosquito population, the current level insecticide efficacy (efficacy typically declines post-deployment as the insecticide, or its matrix, degrades) and the probability of insecticide exposure. Selection is implemented as either a probabilistic process (“polysmooth”, Figure 3 in Hobbs & Hastings (2024)) or by truncation (“polytruncate”, Figure 2 in Hobbs & Hastings (2024)). The response (between generation change) is calculated using the sex-specific Breeder’s equation (Equation 3c, in Hobbs & Hastings (2024)), and cross resistance can be included as a correlated response (Equation 8c, in Hobbs & Hastings (2024)).

### 2.2. Scenario 1: Impact of Pre-Existing Resistance, Insecticide Dosing, and Insecticide Decay

For scenario 1, the pyrethroid partner is given one of four initial resistances (0, 10, 50 and 80% measured as bioassay survival), to cover a range of field realistic resistance levels. The initial resistance for the novel insecticide is 0% bioassay survival, indicating the population is susceptible to the novel insecticide partner.

Mixtures were modelled considering five dosing permutations and running both insecticides (pyrethroid and novel) as full-dose (FD) monotherapies (Table 1). Reduced dose mixtures were modelled assuming a half-dose (HD) retains 50% efficacy (reasonable worst-case scenario) or 75% efficacy (reasonable best-case scenario) of the efficacy of the full dose. The HD_FD and FD_HD simulations (Table 1) are conducted only for a half-dose retaining 50% efficacy as these dosing strategies are used to highlight complexities in IRM evaluation interpretation which arise when insecticides are deployed at unmatched efficacies.

**Table 1:** Scenario Overview for Part 1: Impact of Pre-Existing Resistance, Insecticide Dosing, and Insecticide Decay.

|  |  |  |  |  |  |  |  |  |  |  |
| --- | --- | --- | --- | --- | --- | --- | --- | --- | --- | --- |
| Scenario Description: Level of Resistance |  | Scenario Description: Insecticide Dosing |  |  | Scenario Description: Insecticide Decay |  |  | Scenario Description: Ecological/Biological Parameter Space |  |  |
| Start Polygenic Resistance Score, $\bar{z}_{Pyrethroid}$ | Corresponding Start Bioassay Survival, $K_{Pyrethroid}^B$ (%) | Dosing Scenario ID | Dosing Novel, $\omega_0^{Novel}$ | Dosing Pyrethroid, $\omega_0^{Pyrethroid}$ | Base Decay Rate Novel, $\delta_b^{Novel}$ | Base Decay Rate Pyrethroid, $\delta_b^{Pyrethroid}$ | Decay Threshold Generation, $\tau_b$ | Parameter | Value Range | |
| 0 | 0 | FD_FD | 1 | 1 | 0.005 | 0.005 | 10 | Heritability, $h^2$ | Uniform(0.05 – 0.3) | |
| | | FD_HD | 1 | 0.5 | 0.015 | 0.005 | | Female Exposure, $x$ | Uniform(0.4 – 0.9) | |
| 100 | 10 | HD_FD | 0.5 | 1 | 0.025 | 0.005 | 15 | Male Exposure as a proportion of female exposure, $m$ | Uniform(0 – 1) | |
| | | HD_HD_50 % | 0.5 | 0.5 | 0.005 | 0.015 | | Coverage, $C$ | Uniform(0.5 – 0.9) | |
| 900 | 50 | HD_HD_75 % | 0.75 | 0.75 | 0.015 | 0.015 | 20 | Dispersal, $\theta$ | Uniform(0.1 – 0.9) | |
| 3600 | 80 | Novel Monotherapy | 1 | 0 | 0.025 | 0.015 | | Cross Resistance, $\alpha_{IJ}$ | 0 | |
| | | | | | 0.005 | 0.025 | | Female Fitness Cost, $\phi^{\text{♀}}$ | 0 | |
| | | Pyrethroid Monotherapy | 0 | 1 | 0.015 | 0.025 | | Male Fitness Cost, $\phi^{\text{♂}}$ | 0 | |
|  |  |  |  |  | 0.025 | 0.025 |  |  |  |  |

The efficacy of the insecticides (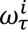 and 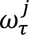) in the mixture are allowed to decay during each deployment interval of 30 mosquito generations (∼ 3 years), which is the standard deployment interval for LLINs. Insecticide efficacy decay is implemented as a two-stage process (Figure 1). During the first stage the insecticides decay slower (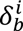 and 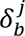). Once a deployment longevity threshold is reached (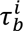 and 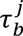, in generations) the insecticides have rapid decay rates (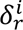 and 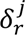). The values of 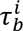 and 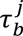 for each insecticide in the mixture are assumed to be the same. For LLINs this may be due to the physical degradation of the net material such that mosquitoes are unlikely to contact the LLIN as so affect each insecticide equally. Details of the implementation of insecticide efficacy decay is detailed in equations 2d(i) and 2d(ii) in Hobbs & Hastings (2024). Details of the estimation of the insecticide efficacy decay rate for standard (pyrethroid-only) LLINs is described in Hobbs & Hastings (2024a). s

**Figure 1.**
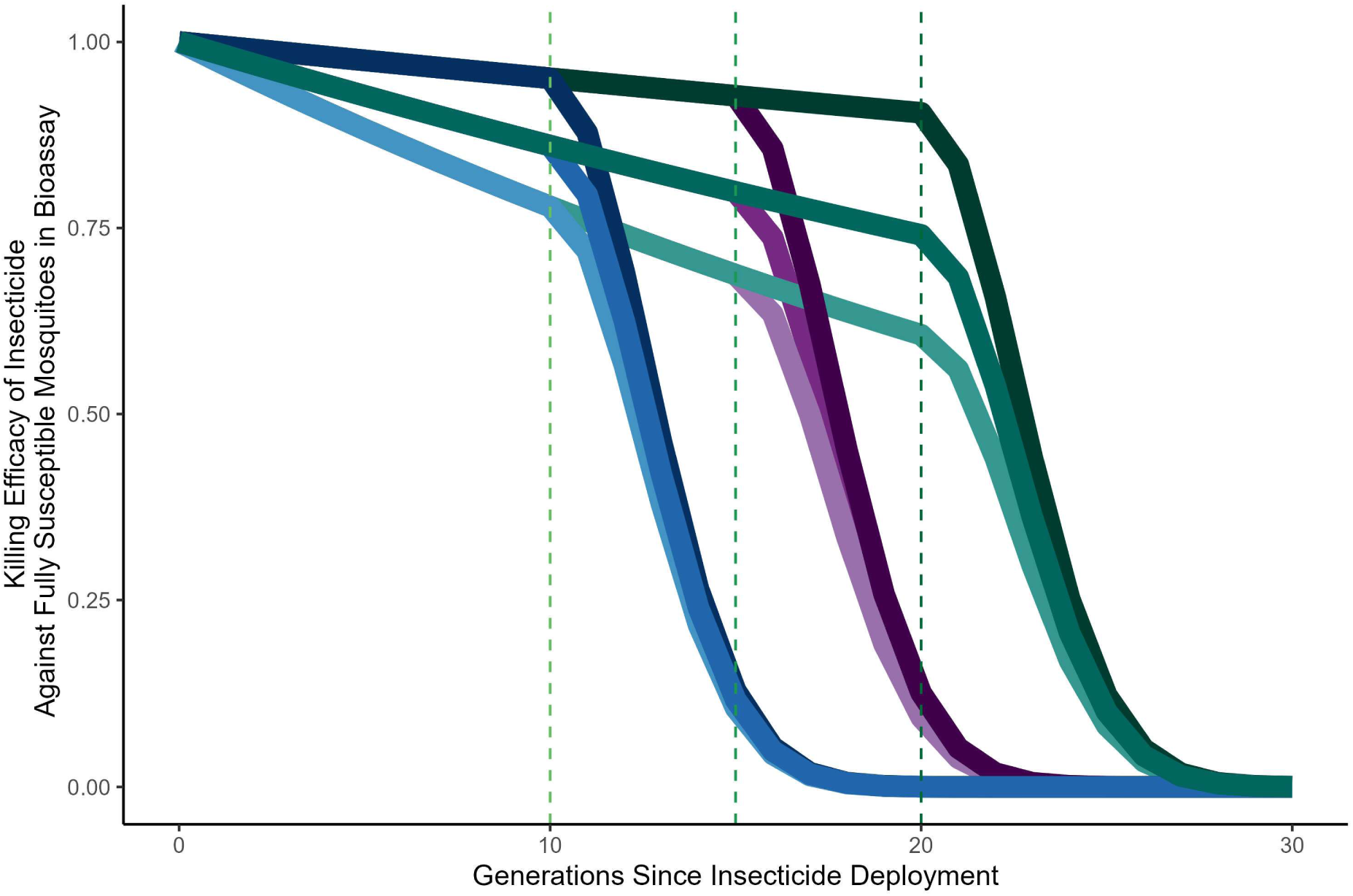
Visualisation of Insecticide Decay Rates. This a visualisation of how insecticides can be expected to decay throughout a simulation. Insecticides are expected to decay slowly on initial deployment, before reaching a threshold time (the dashed lines at 10, 15 and 20 generations) after which the effectiveness of the product, in terms of insecticidal efficacy, rapidly decays due in part to chemistry and the physical integrity of the net.

Latin hyperspace sampling (Carnell, 2020) was conducted to generate a sample of 2500 biological parameter sets, sampling within uniform distributions (Table 1). To simplify interpretation, both insecticide resistance traits (𝐼 and 𝐽) have the same heritability. Cross resistance and fitness costs were not included to simplify scenario interpretation. The issue of cross resistance and insecticide dosing is explored in Scenario 2.

The overview of the simulations for Scenario 1 are given in Table 1, such that all permutations of resistance, dosing and decay are run for each biological parameter set. These were compared against both the novel insecticide being run as a full dose monotherapy for the entirety of the simulation and the pyrethroid insecticide being deployed as a full dose monotherapy. When deployed as monotherapies the insecticides had the same base decay rates and deployment longevity threshold as when the insecticides were deployed in mixture. These simulations were run assuming truncation (“polytruncate”) and probabilistic (“polysmooth”) selection using the same parameter sets allowing for direct comparisons. The standard deviation (𝜎_r_) of the population mean PRS (*z̅_I_*) increases with the value of the mean PRS (Hobbs & Hastings, 2024b), such that the impact of the standard deviation is constant regardless of the magnitude of *z̅_I_*.

Mixtures were compared against pyrethroid monotherapies and novel insecticide monotherapies over a timeframe of 200 generations (∼20 years) on their ability to slow the rate of evolution, defined as the average bioassay change per generation. The rate of evolution for when an insecticide is deployed in monotherapy (𝑀𝑜𝑛𝑜𝑅𝑎𝑡𝑒_i_) is therefore:

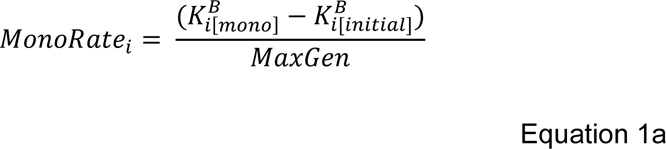

Where 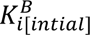 is the initial resistance to the insecticide at the start of the simulation measured as bioassay survival. The subscript 𝑖 denotes the insecticide and is either the novel insecticide or the pyrethroid insecticide. 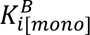 is the bioassay survival to the insecticide at the end of the simulation when the insecticide is deployed as a monotherapy. 𝑀𝑎𝑥𝐺𝑒𝑛 is the number of generations the simulations were run for, being 200 generations in the presented simulations. The rate of evolution for when the insecticide is deployed in mixture (𝑚𝑖𝑥𝑡𝑢𝑟𝑒 𝑟𝑎𝑡𝑒_i_) is therefore:

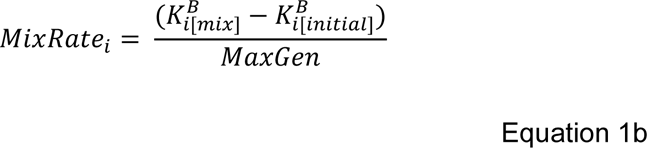

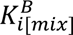 is the bioassay survival to the insecticide at the end of the simulation when the insecticide is deployed in a mixture. The percentage change in the rate of evolution when the insecticide is deployed in mixture versus being deployed as a monotherapy is therefore calculated as:

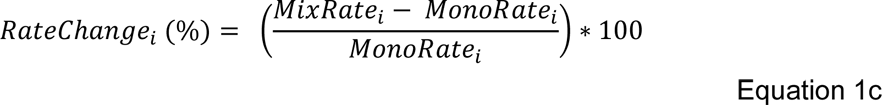

These are calculated separately for the novel insecticide and pyrethroid insecticide, such that the final outcomes are 𝑅𝑎𝑡𝑒𝐶ℎ𝑎𝑛𝑔𝑒_novel_ and 𝑅𝑎𝑡𝑒𝐶ℎ𝑎𝑛𝑔𝑒_pyrethroid_.

The analysis first looks at the broad global level conclusions of the impact of the initial resistance to the pyrethroid, dosing strategy, and insecticide decay separately. The factors of initial resistance to the pyrethroid, dosing strategy, and insecticide decay are then combined to identify more specific trends. Findings are reported as histograms of the rate changes. This primary analysis is conducted grouping “polysmooth” and “polytruncate” together.

Sensitivity analysis was conducted first by separating the “polysmooth” and “polytruncate” simulations and re-conducting the primary analysis to understand the influence of the assumption of the process of selection (Supplement 2). Sensitivity analysis to identify the impact of the biological parameters was conducted by fitting generalised additive models (GAMs), smoothing over the biological parameter space, stratified by dosing strategy and starting resistance to the pyrethroid (Supplement 2). For direct comparisons between “polytruncate” and “polysmooth”, we define the outcome as which mixture strategy was optimal, and then look at the agreement/disagreement between the models (Supplement 3).

### 2.3. Scenario 2: The Impact of Cross Resistance and Dosing

Scenario 2 explores the impact of cross resistance and insecticide dosing on the utility of mixtures for IRM. For simplicity in interpretation insecticide decay is absent, the resistance traits have the same heritability, and both insecticides have an initial starting bioassay survival of 0%.

Cross resistance is included at fixed values of -0.5 to 0.5, at 0.1 intervals (Table 2). Negative values indicate negative cross resistance where resistance to one insecticide increases susceptibility to the other insecticide. Positive values indicate positive cross resistance where resistance to one insecticide increases the resistance to the other insecticide. Three mixtures doses are evaluated i.e., a full-dose mixture (FD_FD) or half-dose mixture assuming the half-dose retains either 50% (HD_HD 50%) or 75% (HD_HD 75%) of the efficacy of the full dose (Table 2).

**Table 2:**
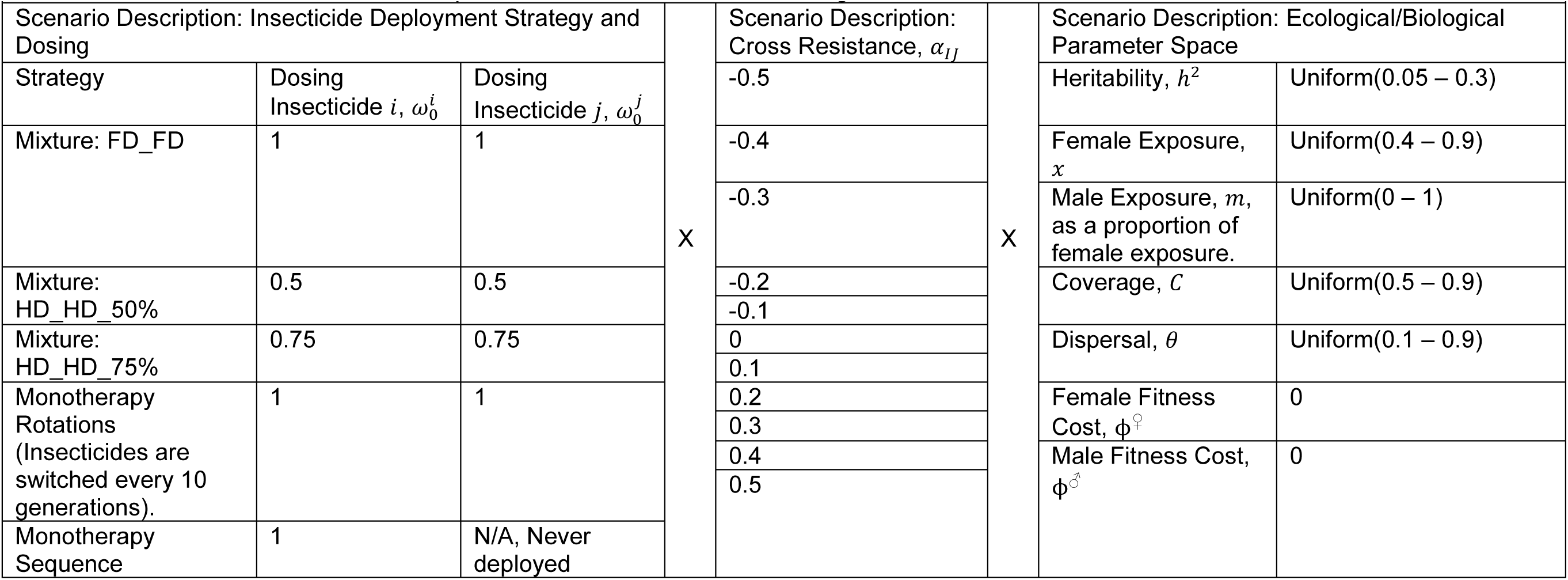
Scenario Overview for Part 2: Impact of Cross Resistance and Dosing.

These three mixture doses were compared against each other and against 1.) continuous monotherapy of one insecticide and 2.) deploying insecticides 𝑖 and 𝑗 in rotation (as monotherapies) with a deployment interval of 10 generations (Table 2). All simulations run for 200 generations (∼20 years). The continuous monotherapy of one insecticide comparator was used to highlight the issues which can arise when withholding insecticides for fear of cross resistance.

All permutations of the cross-resistance values, mixture doses and biological parameter sets were conducted (Table 2). These were compared against insecticide 𝑖 deployed as a monotherapy (with insecticide j never being deployed) and insecticide 𝑖 and insecticide 𝑗 deployed in rotation for all permutations cross-resistance values and biological parameter sets to allow for direct comparisons. This process was conducted assuming truncation (“polytruncate”) and probabilistic (“polysmooth”) selection using the same parameter sets allowing for direct comparisons.

At the end of the simulations (200 generations) the bioassay survival to each insecticide was extracted and the difference between the mixture simulations and the comparators strategies was calculated. The difference in bioassay survival at the end of the simulations is calculate for each insecticide.

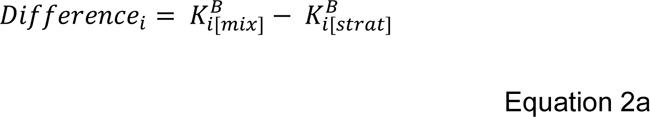

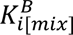 is the bioassay survival to insecticide 𝑖 at the end of the simulation (after 200 generations), when the insecticide is deployed in mixture. 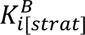 is the bioassay survival to insecticide 𝑖 at the end of the simulation when the insecticide is deployed in any of the comparator strategies (continuous monotherapy, rotation monotherapies, or the alternate mixture doses). This means the FD_FD simulations are compared against each HD_HD (retaining 50 or 75% efficacy) simulation and the two HD_HD strategies are also compared against each other. The difference for insecticide 𝑗 is also calculated:

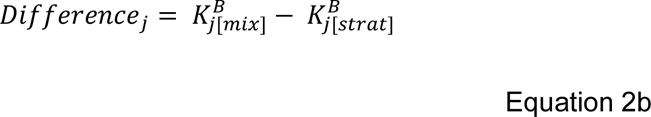

Finally, the total difference in bioassay survival at the end of the simulations is calculated:

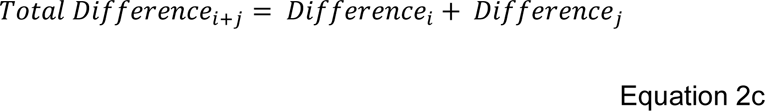

The differences in the bioassay changes are reported and histograms are used to visually assess the differences. This analysis is grouped for “polytruncate” and “polysmooth” and is also conducted separately to understand the impact of the assumption regarding the insecticide selection process (Supplement 2).

Sensitivity analysis for the biological parameters was conducted using GAMs. Comparisons between “polytruncate” and “polysmooth” were made by comparing which dosing strategy performed best and comparing the model agreements.

## 3. Results

### 3.1. Scenario 1: Impact of Pre-Existing Resistance, Dosing and Decay

#### 3.1.1. Scenario 1: Impact of Mixture Dosing

Looking at the impact of the mixture dosing (Figure 2) two global level conclusions can be drawn. First, that full-dose mixtures (FD_FD) perform best to protect both the novel insecticide and the pyrethroid insecticide simultaneously. When looking at the FD_HD and HD_FD dosing strategies, the protection offered to one of the insecticides is at the expense of the other insecticide.

Second, reducing the dose (and therefore the effectiveness) of the insecticides in mixture reduces the benefit of the mixture. Simulations where we allowed the half dose to retain 75% of the full dose efficacy, show this dosing strategy is more effective than when it retains only 50% of the efficacy but is still less effective than the full dose mixtures (Figure 2). This pattern is consistent between “polytruncate” and “polysmooth” (Supplement 2, Figure S2.1).

#### 3.1.2. Scenario 1: Impact of Initial Resistance to Pyrethroid

Next, we examine the impact of the initial resistance to the pyrethroid. Figure 3 shows that as initial resistance to the pyrethroid increases, the benefit of deploying the novel insecticide in mixture with the pyrethroid decreases. In addition to this, as the rate of evolution to the novel insecticide increases with resistance to the pyrethroid insecticide (due to lack of protection), using mixtures to slow the rate of evolution in the pyrethroid also becomes less effective as the resistance to the pyrethroid increases. Once again, “polytruncate” and “polysmooth” gave similar conclusions (Supplement 2, Figure S2.2).

#### 3.1.3. Scenario 1: Impact of Insecticide Decay

Figure 4 shows how mismatches in the rates of insecticide decay of the two insecticides impacts their ability to slow the rates of evolution. Differential decay rates are often highlighted as a concern for mixtures (e.g., Denholm & Rowland, 1992), our results appear to indicate this is less of a concern than previously thought when considering long evolutionary trajectories. The impact of differential decay rates is lower than the impact of the chosen dosing strategy or initial mismatches in the amount of resistance to both insecticides (Figures 2 and 3). Again, “polytruncate” and “polysmooth” models gave similar conclusions (Supplement 1, Figure S1.3).

**Figure 2.**
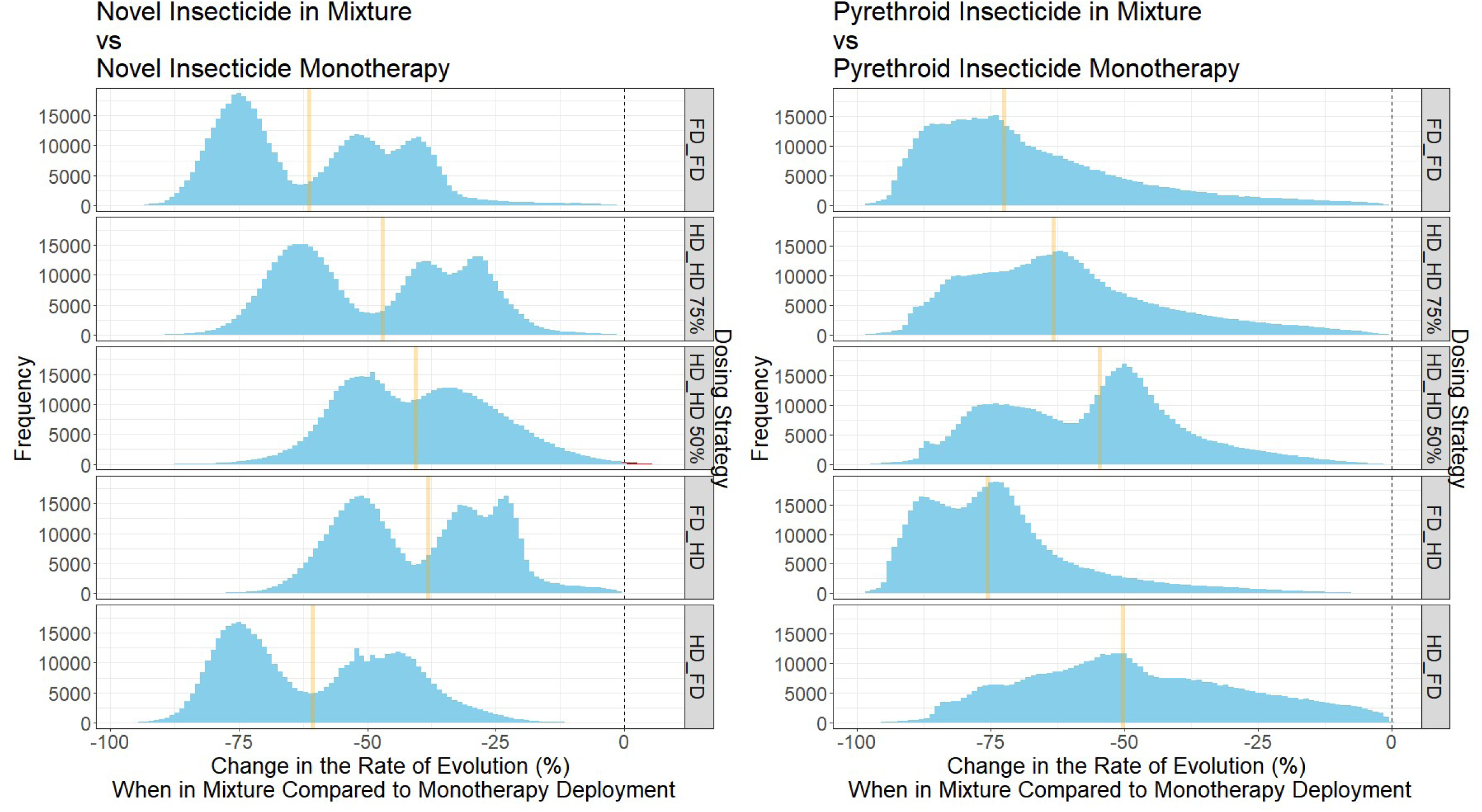
Comparing the Mixture Dosing Strategies versus Monotherapy Deployments Stratified by Dosing Strategy. Left panels is comparing each mixture dosing strategy versus deploying the novel insecticide monotherapy at full dose. The right panel is comparing each mixture dosing strategy versus deploying the pyrethroid insecticide alone. Orange line indicates the median value. Red bars indicate simulations where deploying as mixture was found to be the worse strategy (i.e. with values greater then 0). Dosing strategies are in the form Novel_Pyrethroid, with FD=full dose and HD = half dose. The larger the negative numbers (i.e. to the left on the X-axes), the greater the benefit of deploying the insecticides in the mixture versus monotherapy deployment.

**Figure 3.**
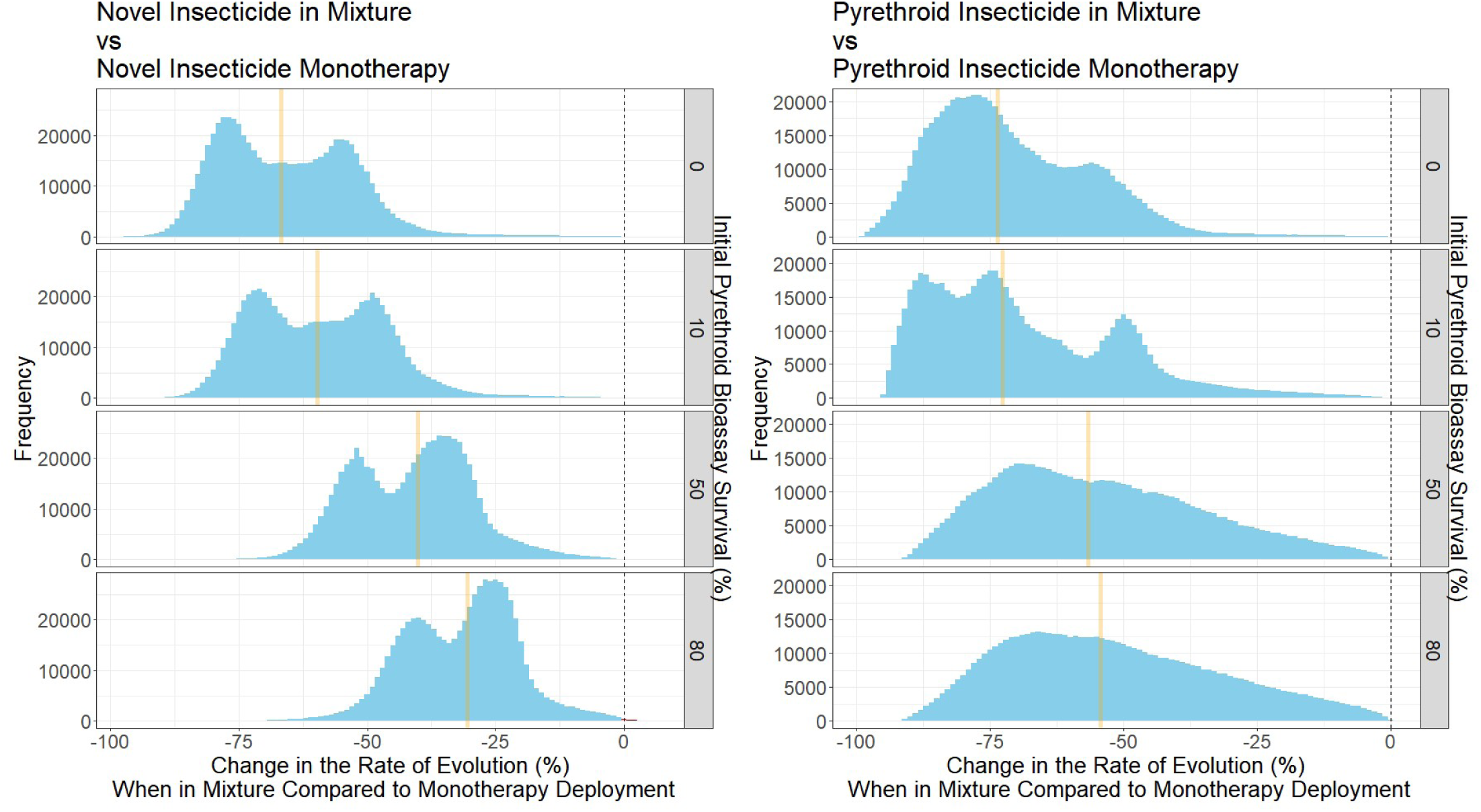
Comparing mixture deployments versus monotherapy deployments stratified by initial bioassay survival to the pyrethroid. The “rate of evolution” is defined as the change in bioassay survival over a 200 generation (∼20 year) time horizon. Left Panel is comparing the mixture dosing strategy versus deploying the novel insecticide monotherapy at full dose. The right panel is comparing the mixture dosing strategy versus deploying the pyrethroid insecticide alone. Orange line indicates the median value. Red bars indicate simulations where deploying as mixture was found to be the worse strategy (i.e. when values are positive). Each panel indicates how much pyrethroid resistance was present at the start of the simulation, measured as bioassay survival (%).

**Figure 4.**
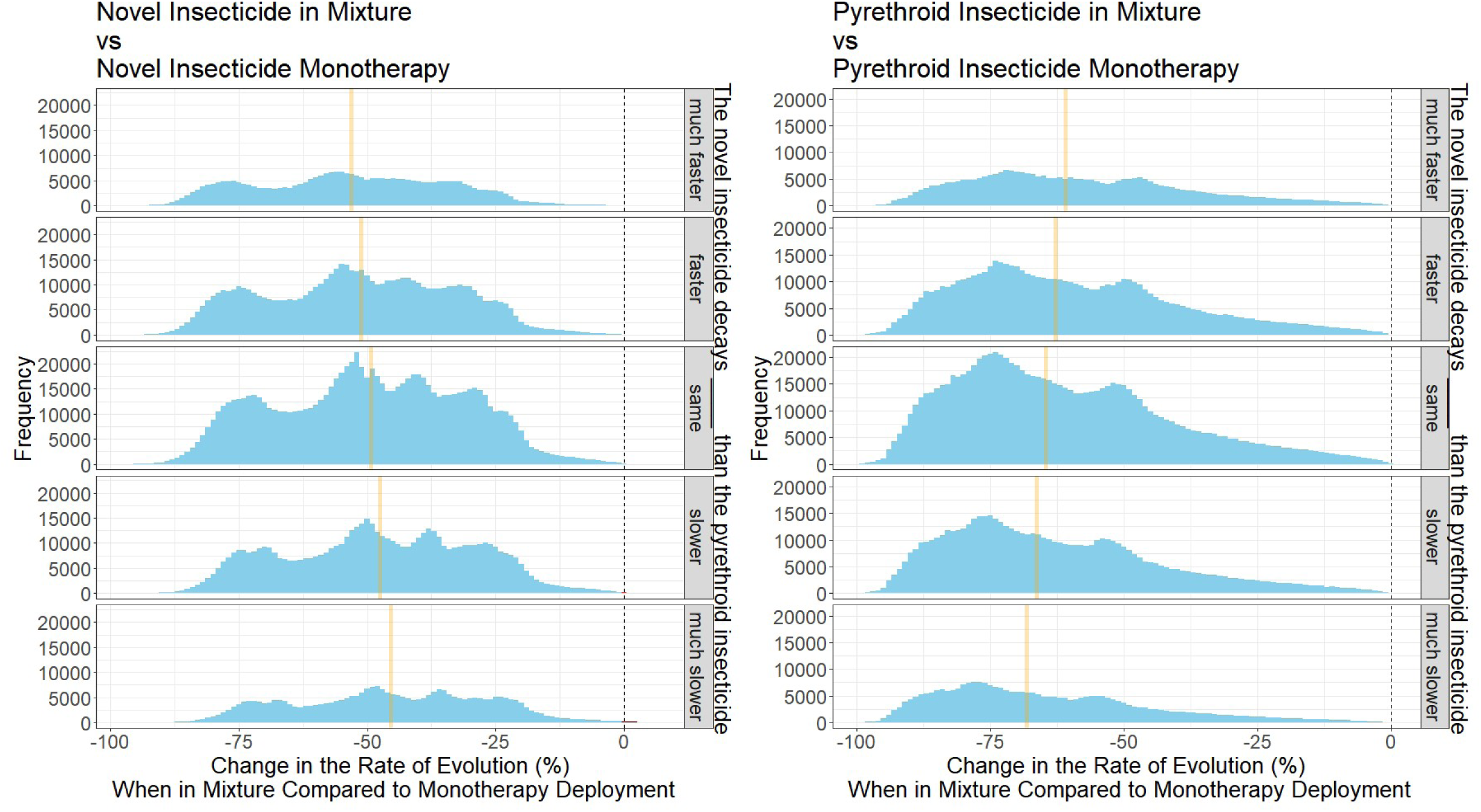
Comparing mixture deployments versus monotherapy deployments stratified by decay rates. Left Panel is comparing each mixture dosing strategy versus deploying the novel insecticide monotherapy at full dose. The right panel is comparing each mixture dosing strategy versus deploying the pyrethroid insecticide alone. Orange line indicates the median value. Red bars indicate simulations where deploying as mixture was found to be the worse strategy. The description at the side of each panel describes how the novel insecticide decays with respect to the pyrethroid, and therefore should be read as: The novel insecticide decays “description” than the pyrethroid insecticide.

#### 3.1.4. Scenario 1: Impact of Interaction of Initial Resistance to Pyrethroid, Mixture Dosing and Insecticide Decay

When looking at how the initial resistance to the pyrethroid, the mixture dosing strategy and decay rates interact (Figure 5) we see that a full dose of both insecticides is the best dosing strategy, with the advantage of slowing the rate of resistance to both the insecticide partners. A half dose of both insecticides was found to be overall the worst dosing strategy (of those evaluated), only reducing the rate of evolution marginally compared to full dose monotherapy deployments. A full dose of the novel insecticide and half dose of the pyrethroid was found to slow the rate of evolution in the pyrethroid, but at the expense of the novel insecticide. And likewise, a half dose of the novel insecticide and full dose of the pyrethroid was found to slow the rate of evolution in the novel insecticide but at the expense of the pyrethroid.

**Figure 5.**
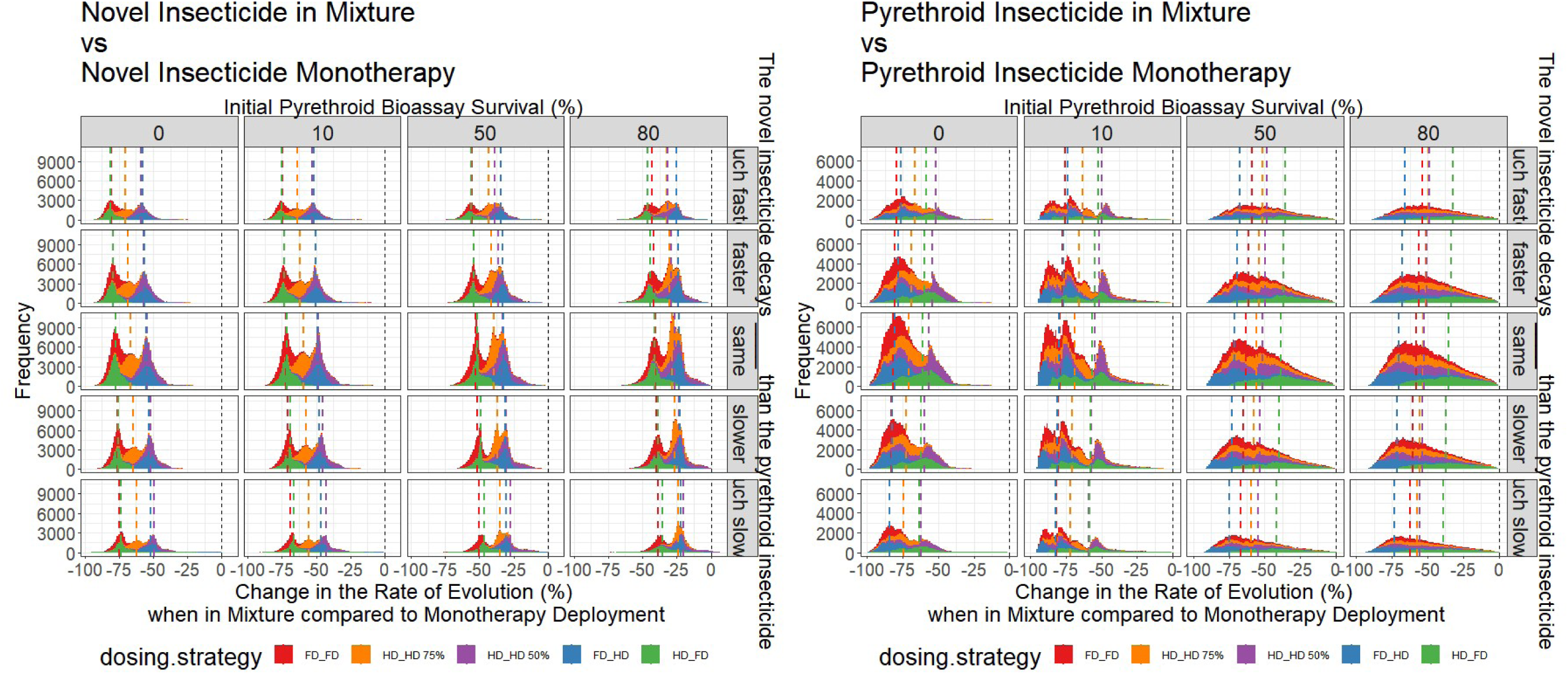
Comparing mixture deployments versus monotherapy deployments stratified by decay rates, initial pyrethroid resistance and mixture dosing. Colours are the dosing strategies. Red is full dose both insecticides. Purple is half dose of both insecticides. Green is full dose of the novel insecticide and a half dose of the pyrethroid. Blue is the half dose of the novel insecticide and a full dose of the pyrethroid. Here, we are assuming a half dose retains 50% of the efficacy of a full dose. Stratification left to right is pre-existing resistance to the pyrethroid, and stratified top to bottom is the differences in decay rates. Left panels are the novel insecticide and the right panels are the pyrethroid insecticide.

When comparing the mixture dosing strategies directly against one another, we see that the full dose mixture is best, followed by the mixture which retains 75% of efficacy, with the mixture retaining 50% of the efficacy performing generally worst. However, there are occasions when this pattern did not hold (Figure 6), which can be explained by the sensitivity analysis.

**Figure 6.**
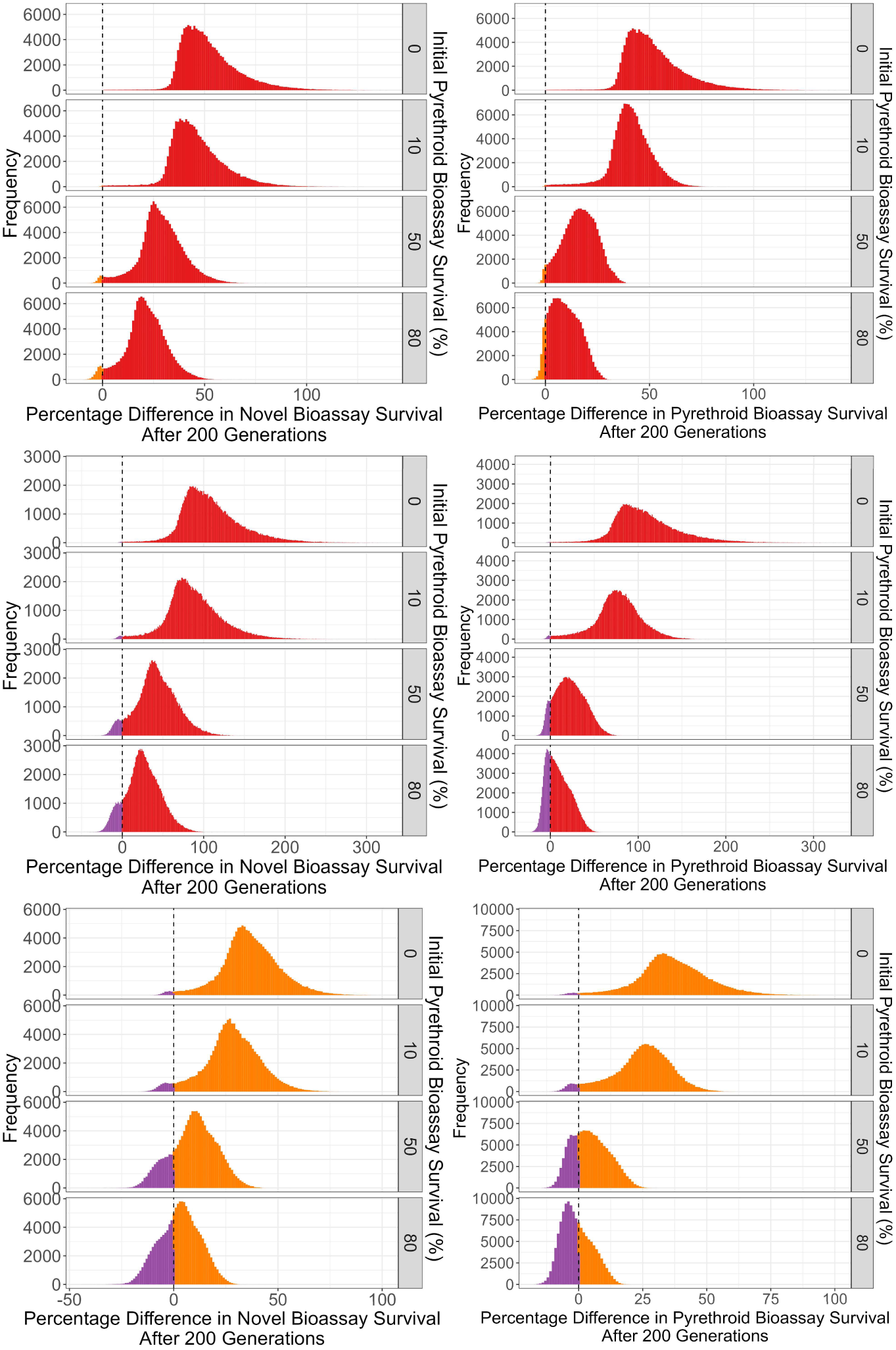
Direct comparisons between mixture dosing strategies. The percentage difference in the rates of evolution (measured as the change in bioassay after 200 generations) to each insecticide partner was assessed. Left plots = novel insecticide, right plots = pyrethroid insecticide. The colours indicate which dosing strategy performed better in the direct comparisons, where: Red = FD_FD, Purple = HD_HD retains 50% efficacy, Orange = HD_HD retains 75% efficacy.

#### 3.1.5. Scenario 1: Interaction with Biological Parameters

Biological variables outside of operational control vary between locations. It is therefore important to understand how these variables may impact IRM. Figure 7 reveals insight into the conditions required for the benefit of FD_FD mixtures above half-dose mixtures. First, is when there is substantial initial resistance to the pyrethroid (>50% bioassay survival) then at high levels of female insecticide exposure (>0.8) then the HD_HD mixture looks to be the better dosing strategy when looking to protect the novel insecticide when compared to the FD_FD mixture. At very high levels of mosquito exposure, only the most resistant individuals would survive and constitute the parents of the next generation, that is there would be fewer less resistant individuals managing to avoid the insecticide encounter. This does support the long-held hypothesis that for mixtures to be effective as a IRM strategy a proportion of the population must escape insecticide exposure to have a more susceptible portion of the population to dilute the overall resistance of the population. However high coverages of high dose mixtures would be expected to provide better control (Madgwick & Kanitz, 2022a). Additional sensitivity analysis GAMs can be found in Supplement 2.

**Figure 7.**
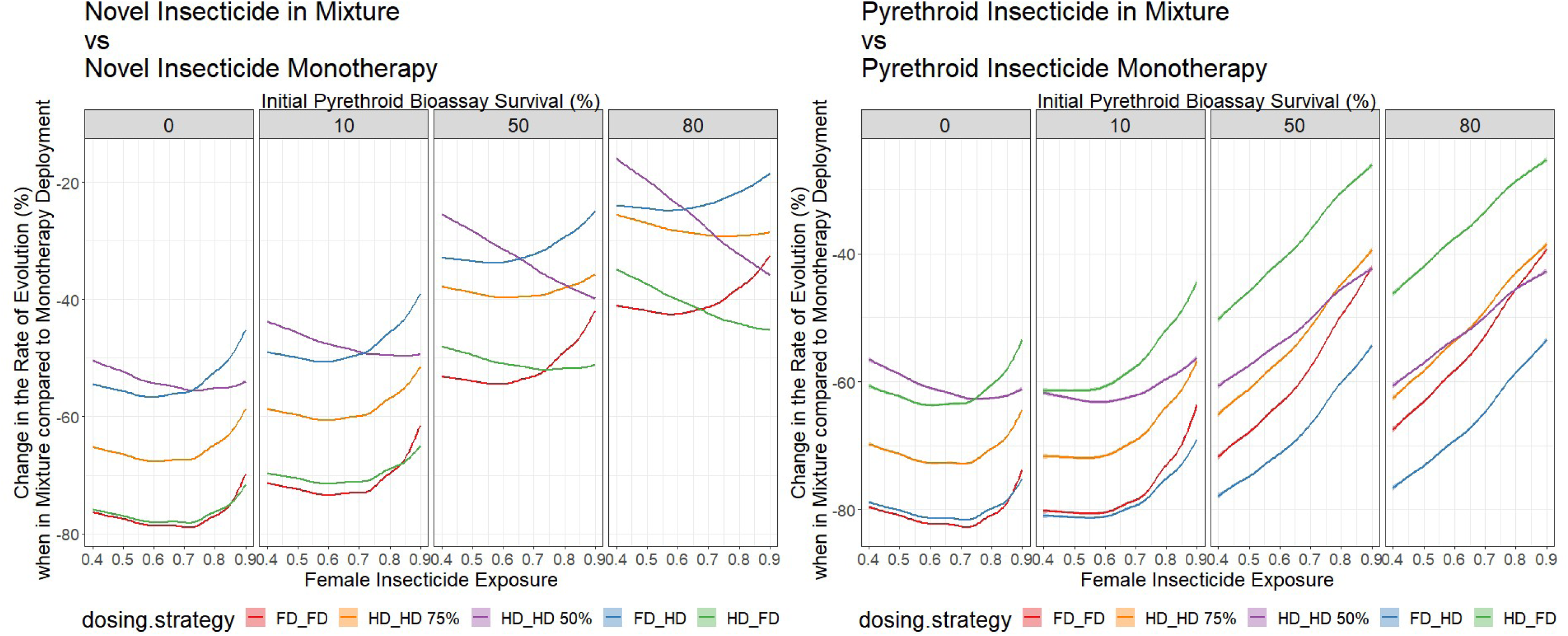
Generalised Additive Model: Change in Rate of Evolution as a function of Female Exposure. Interpretation is that the lower the line, the better the dosing strategy is for slowing the rate of evolution. Left panels are for mixtures compared to monotherapy deployment of the novel insecticide, right panels are for mixtures compared to monotherapy deployment of the pyrethroid.

From Scenario 1 the following messages key messages are highlighted:

1. Increasing levels of initial resistance to the pyrethroid reduces the long-term benefit of insecticides being in mixture (Figures 3 and 5).
2. Full dose mixtures are generally more effective than half dose mixtures in slowing the rate of evolution (Figures 2, 5, 6, 7).
3. Insecticide decay mismatches appear to be less of an issue than previously thought over the whole operational lifespan of the mixture i.e., variability in initial levels of resistance and/or dosing play a more noticeable role (Figure 4).

### 3.2. Scenario 2: Impact Cross Resistance and Insecticide Dosing for Mixtures

#### 3.2.1. Scenario 2: Cross Resistance and Insecticide Dosing: Implication of Withholding Insecticides

The deployment of insecticide 𝑖 and 𝑗 in mixture was compared against the deployment of insecticide 𝑖 only (continuous monotherapy) to simulate the impact of withholding insecticide 𝑗 from deployment. For insecticide 𝑖, we can see the mixture (regardless of the mixture dosing or level of cross resistance) performed better at slowing resistance to insecticide 𝑖. This would of course be expected, as the mixtures are being compared to the deployment of only one insecticide (Figure 8).

**Figure 8.**
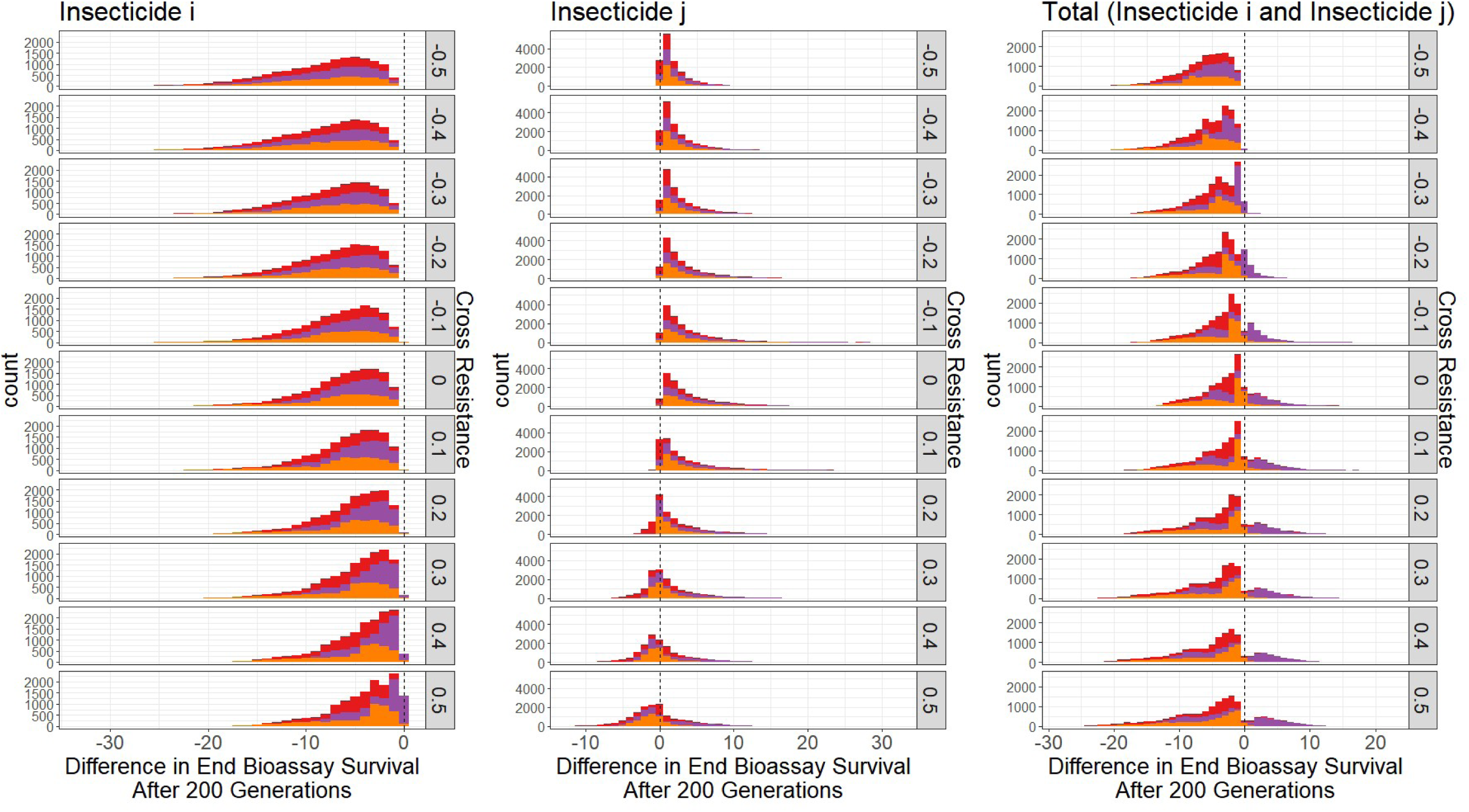
Change in bioassay survivals (absolute) when deployed as a mixture (𝒊 and 𝒋) versus a continuous monotherapy of insecticide 𝒊 after 200 generations with cross resistance. Left panel is the change between the monotherapy deployed insecticide and the insecticide when in mixture. Middle panel is the change in the change between insecticide 𝑗 (never deployed) and when deployed in mixture. Right panel is the total change in bioassay survival (sum of end bioassay survivals). Red = FD_FD, purple = HD_HD retains 50%, orange = HD_HD retains 75%. Rows are the degree of cross resistance between the insecticides.

For insecticide 𝑗 however, resistance developed faster in the mixture when there was negative or no cross resistance, this is expected as in the continuous monotherapy of insecticide 𝑖 there is never any selection on insecticide 𝑗. However, when there is positive cross resistance, resistance develops slower to insecticide 𝑗 when in mixture than if insecticide 𝑗 had not been deployed (Figure 8). This highlights an important concern regarding “holding back” insecticides, as by the time they are needed to be used it is not unrealistic that cross resistance may have already led to substantial resistance to those insecticides (Figure 8).

When considering the “total amount of resistance” (the sum of the bioassay survivals to each insecticide), we see HD_HD retaining 50% efficacy performs worse than if just insecticide 𝑖 was deployed alone, while FD_FD and HD_HD retains 75% efficacy perform better than the continuous monotherapy deployment of insecticide 𝑖 (Figure 8).

#### 3.2.2. Scenario 2: Cross Resistance and Insecticide Dosing: Mixtures versus Rotations

When comparing mixtures versus rotations in the presence of cross resistance (Figure 9) we see with positive cross resistance higher dose mixtures perform better than rotation. The lower the insecticide dose in the mixture the worse the performance, such that HD_HD (50% efficacy) mixture often performs worse than deploying each insecticide as full-dose monotherapies in rotation. When there is negative cross resistance, the full-dose mixture still performs best. However, at extreme levels of negative cross resistance the benefit of the mixtures against rotations is lower because the rate of selection is low regardless of the strategy.

**Figure 9.**
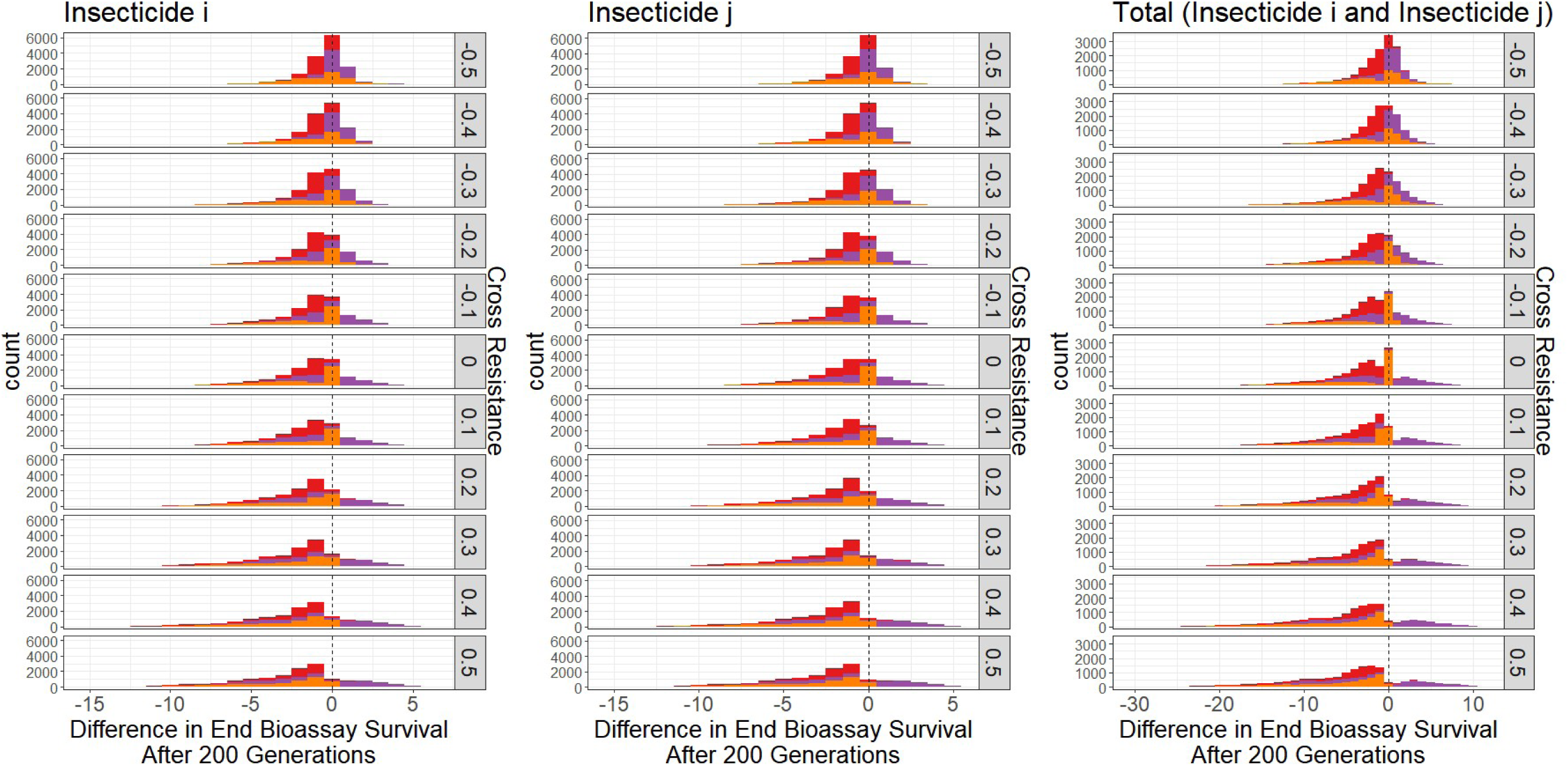
Impact of Cross Resistance on the Efficacy of Mixtures versus Rotations to slow the rate of evolution of insecticide resistance. Graphs are stratified by the amount of cross resistance between the two insecticides. Colours of the bars in indicate the dosing strategy of the mixture, where Red = both insecticides at full dose, orange = both insecticides at half dose and retain 75% of the full dose efficacy, and purple = both insecticides at half dose and retain 50% of their full dose efficacy.

When considering simulations where there was no cross resistance between insecticides, the HD_HD (50% efficacy) performed worse for each individual insecticide ending with a higher bioassay survival. This is an interesting finding as the HD_HD (50% efficacy) contains the same amount of total efficacy as a single full dose insecticide. This indicates the benefit of mixtures is not necessarily from mixing but from mixtures having a greater total amount of insecticide (Figure 9).

From Scenario 2, we would like to highlight the following messages:

1. Withholding insecticides from deployment can mean that substantial resistance evolves because of cross resistance (Figure 8).
2. Full dose mixtures appear most effective, even in the presence of cross resistance (Figure 9).
3. Half dose mixtures perform poorly in the presence of cross resistance (Figure 9).

### 3.3. Qualitative comparison between “polytruncate” and “polysmooth”

We should note that the qualitative outcome agreement (i.e., which dosing strategy was best) showed a very good level of agreement between “polysmooth” and “polytruncate”. A small subset of simulations disagreed. The choice of a default mixture strategy, which would generally perform best remained the full dose mixture regardless of the assumed selection process. See Supplement 3 for details.

## 4. Discussion

Using dynamic models of insecticide selection assuming polygenic resistance (Hobbs & Hastings, 2024b), we have explored the conditions which may be beneficial/detrimental to the use of mixtures as part of an IRM strategy. Considering the numerous conclusions which can be reached from our presented results (a headline summary of which can be found in Table 3) we focus our discussion on what we consider to be the key take-home messages:

1. Full dose mixtures perform best for slowing the rate of evolution (Figures 2, 5, 7, 8 and 9).
2. Half dose mixtures can be worse than deploying the equivalent insecticides individually as a rotation (Figure 9).
3. Mixtures are most beneficial from an IRM perspective when there is limited resistance to either insecticide (Figures 3, 5 and 6).
4. Full dose mixtures may be able to overcome cross resistance (Figures 8 and 9).

**Table 3:** Summary Table of Figures.

| **Table 3: Summary Table of Figures** |  |
| --- | --- |
| In this table we highlight the key messages for the figures presented to aid in interpretation. |  |
| Figure | Summary/Interpretation |
| 1 | • During an insecticide deployment, insecticide will decay.<br>• This will depend on both insecticide chemistry and environmental conditions. |
| 2 | • Full-dose mixture most effective.<br>• Half-dose mixture (retaining 50%) least effective.<br>• Unequal doses protect one insecticide at the expense of the other. |
| 3 | • As resistance to pyrethroid increases, benefit of mixtures decreases. |
| 4 | • Differential decay rates have lower impact than issues surrounding dosing and initial levels of resistance over the long term. |
| 5 | • Increasing resistance decreases benefit of being in mixture<br>• Reduced dosage less effective (for IRM) than higher doses.<br>• Differential decay rates have minimal impact over the long term. |
| 6 | • Effective mixtures require some of the population to escape selection. |
| 7 | • Full-dose mixtures outperform half dose mixtures in direct comparisons.<br>• Reduced dose mixtures need to retain high efficacy. |
| 8 | • Deploying HD_HD mixtures can lead to situations where there is more resistance evolution than if only one insecticide is deployed.<br>• Deploying HD_HD mixtures worse than deploying FD_FD mixtures. If HD_HD retains only 50% efficacy, can be worse than just deploying a monotherapy insecticide. |
| 9 | • HD_HD retaining 50% efficacy can be worse than deploying each insecticide in rotation as a monotherapy.<br>• FD_FD performs better (or equal) to rotations regardless of cross resistance.<br>• When there is very negative cross resistance all strategies perform near equivalently. |

### 4.1. Full dose mixtures perform best for slowing the rate of evolution

As highlighted in Figures 2, 5, 8 and 9, mixtures containing a full dose of both insecticides have been found to best slow the rate of evolution to both insecticide partners, and this was consistent regardless of the amount of resistance present to the insecticide partners and the amount of cross resistance between the two insecticides. However, our parameter sensitivity analysis has highlighted the superiority of full dose mixtures requires a portion of mosquito population to escape selection (Figure 7), which is consistent with monogenic models (Rex Consortium et al., 2013). This requirement would seem likely as LLIN and/or IRS coverage rarely exceeds 70% (Bhatt et al., 2015). The endophagic and/or endophilic behaviours are also unlikely to be exclusive, with *Anopheles* species showing plasticity in host preferences (Takken & Verhulst, 2013). These modelling results are at odds with current mixture products in development and deployment, which appear to be reducing the dosing of at least one of the insecticide partners. This finding of full-dose mixtures being most effective is consistent with the monogenic modelling literature (Levick et al., 2017; Madgwick & Kanitz, 2022b). We should note that the qualitative outcome agreement (i.e., which dosing strategy was best) showed a very good level of agreement between “polysmooth” and “polytruncate” (Supplement 3), with only a small subset of simulations disagreeing. The choice of a default mixture dosing strategy, which on average would perform best, remained the full-dose mixture for either mechanism of selection.

### 4.2. Half-dose mixtures may be worse than deploying the equivalent insecticides individually as a rotation

The Rex Consortium highlighted that reduced dosage mixtures were infrequently evaluated (Rex Consortium, 2013). Our findings indicate deploying two insecticides as monotherapies over time (in rotation) was as effective for IRM strategy as deploying two insecticides as half-dose mixtures (Figure 9). Here, we are assuming there is the same “total amount” of insecticide in a half-dose mixture as in a full-dose monotherapy deployment. We therefore explored whether the effect of mixing is advantageous and the benefit not just there being double the amount of insecticide in full-dose mixtures which is often believed to inflate the efficacy of mixtures (Rex Consortium, 2013). From this we can perhaps consider the benefit of full dose mixtures is less to do with the increased chemical diversity, and more to do with the increased killing potential of having two insecticides at full-dose. It is most likely that the effect is already as postulated by monogenic models i.e., that high levels of killing efficacy are required in both insecticides as they provide effective mutual protection. In effect there is little advantage in an insect being resistant to insecticide 𝑖 if it is reliably killed by insecticide j in the mixture Comparing this result against the monogenic literature is problematic, due to half-dose mixtures being rarely explored in monogenic systems, despite being highlighted as a potential issue as early as the 1980s (Curtis, 1985). Levick et al (2017) did quantify the impact of reduced efficacy of insecticides in a mixture, as it likely to occur when half-doses are used, and found that this severely comprised the advantages of a mixture, to the extent that they may become counter-productive and insecticides would be better deployed alone. This conclusion is in line with that obtained here from our polygenic model. Helps et al. (2017) found that the equivalent of a half-dose mixture performed best, when compared against full-dose mixtures and rotations, in contrast to our results. However, whether this is due to modelling as a monogenic trait or using a different outcome for evaluating the effectiveness of the IRM strategy (Helps et al. (2017), used time until mortality by the insecticide fell below 50%) is unclear. The benefit of additional kill, and therefore additional control, which is expected from full-dose mixtures is frequently not considered when modelling IRM (Madgwick & Kanitz, 2022a).

### 4.3 Mixtures perform are most beneficial from an IRM perspective when there is limited resistance to either insecticide

The benefit of deploying two insecticides in a mixture is greatest when the target mosquito is susceptible to both the insecticides (Figures 2, 5 and 6), and this effect was consistent regardless of the permutation of parameter and mechanism of selection. This finding of mixtures being most beneficial when resistance to both insecticides is low is consistent with the monogenic modelling literature (Rex Consortium, 2013). Unfortunately, situations where mosquito populations are fully susceptible to both the insecticides which are to be used in the mixture is also the situations where IRM is least urgently needed and implemented. The implementation of IRM strategies is most perceived to be needed in situations where resistance has already evolved and is an operational problem. We would therefore re-emphasise the need to implement IRM strategies pre-emptively rather than react once resistance reaches concerning levels, as has been done numerous times (e.g. WHO, 2012). Trials of the next-generation mixture LLINs are also indicating that they appear to better at slowing the rate of pyrethroid resistance evolution than standard pyrethroid-only LLINs (Messenger et al., 2023), however as this trial did not survey all clusters, it is unclear how generalisable this finding is, and how the IRM capability of next-generation mixture LLIN performs over longer time horizons.

### 4.4 Full dose mixtures may be able to overcome cross resistance

We should take special care with our results and subsequent conclusions with regards to cross resistance. Our results indicate in a hypothetical situation where only two insecticides are available and there is positive cross resistance between them, the best course of action would be to combine both these insecticides at full-dose into a single mixture formulation (Figures 8 and 9). It may well be that positive cross resistance is more likely to occur between insecticides than negative cross resistance. For example, insecticides deployed as LLINs or IRS require penetrating the mosquito cuticle, and therefore any cuticular resistance is likely to affect all insecticides deployed in this manner.

The finding that full dose mixtures are more effective (when compared to monotherapy rotations) in the presence of cross resistance therefore increases its utility as a default “go-to” IRM strategy when there are no local field data regarding resistance phenotypes or mechanisms of resistance. Few monogenic models have included cross resistance but these findings are in agreement with a monogenic model of transgenic crops with pyramiding two toxins (i.e., acting as a mixture) remained more effective than sequential deployment with positive cross resistance (Roush, 1998).

### 4.5. Operational Implications for Mixtures for Vector Control

IRM is only one aspect to consider when deploying insecticides for vector control. Considerations must also be made regarding disease control and economics. How much better does a mixture need to be to be viable economically? An important consideration of any IRM plan is also an evaluation of its economic benefits. For insecticide mixtures it may be reasonably expected that developing a mixture product is more costly than a single insecticide product. Mixture formulations will need to be commercially viable for a longer period to justify the increased economic costs associated with development and manufacturing. Half dose mixtures do not appear to decrease the rate of evolution much compared to a full dose novel only deployment. The economic evaluation of IRM is also becoming a more important consideration (WHO, 2021).

As developing mixtures is a complex and expensive challenge, especially when compared against developing single insecticide formulations. A question then arises as to whether fine scale spatial deployments of single insecticide formulations can be used to obtain the effect of a mixture as the mosquitoes encounter both insecticides throughout their own lifespan. There are two potential ways in which this may be conducted. First is the combination of LLIN and IRS within the same household. The use of IRS as part of an IRM has the benefit of further expanding the insecticide arsenal (as more insecticides are available for use as IRS than on LLINs) and allowing shorter timeframes between deployments (IRS is replenished more frequently than LLINs), thereby allowing swifter replacement of failed or failing insecticides. The second strategy to consider is micro-mosaics (Jones et al., 2023). Micro-mosaics involve different households in the same village receiving different brands of LLIN which can then contain different insecticides. Future modelling work will look at the role of combinations and micro-mosaics as possible IRM strategies, and whether the effect of a full-dose mixture can be obtained through fine-scale spatial deployments.

### 4.6. Conclusions

Our results highlight that the use of full-dose mixtures is likely the IRM strategy with the lowest chance of being the worst strategy to implement. Implementing full-dose mixtures pre-emptively will also prolong the utility of this strategy, rather than waiting until resistance to the pyrethroid a critical threshold after which the switch is made.

## Author Contributions

- NPH: Data Curation, Software, Formal Analysis, Investigation, Methodology, Writing – Original Draft Preparation
- IMH: Conceptualisation, Investigation, Methodology, Writing – Review & Editing, Supervision

## Funding Information

NPH was funded by a Medical Research Council – Doctoral Training Partnership grant (2269329). The funder had no role in the in the study design, interpretation of data, writing of the paper or decision to publish.

## Conflict of Interest Statement

The authors declare no conflict of interest.

## Data Availability Statement

Model code for the running and analysis of simulations is available from the github respository: https://github.com/NeilHobbs/polytruncate and https://github.com/NeilHobbs/polysmooth and the corresponding author upon request.

## Acknowledgements

We would like to acknowledge the advice of David Weetman for help grounding the model in biological and operational relevance. We would also like to acknowledge the Vector Informatics and Genomics group at the Liverpool School of Tropical Medicine for their general critiques of the model methodology.

## Notes

### Competing Interest Statement

The authors have declared no competing interest.

