## Supplement 1 for "Exploring operational requirements of mixtures for insecticide resistance management in public health using a mathematical model assuming polygenic resistance"

### Supplement 1: Sensitivity Analysis

An important consideration in our analysis is the implication of how the selection process was implemented. We model selection as happening as a probabilistic process (“polysmooth”) or as threshold selection process (“polytruncate”). The question is how these two implementations differ and/or agree with each other. We therefore present the graphs presented in the main manuscript again, this time separated by the model (“polysmooth” or “polytruncate”). We also provide a summary of the similarities and differences for these plots in Table S1 in this supplement also.

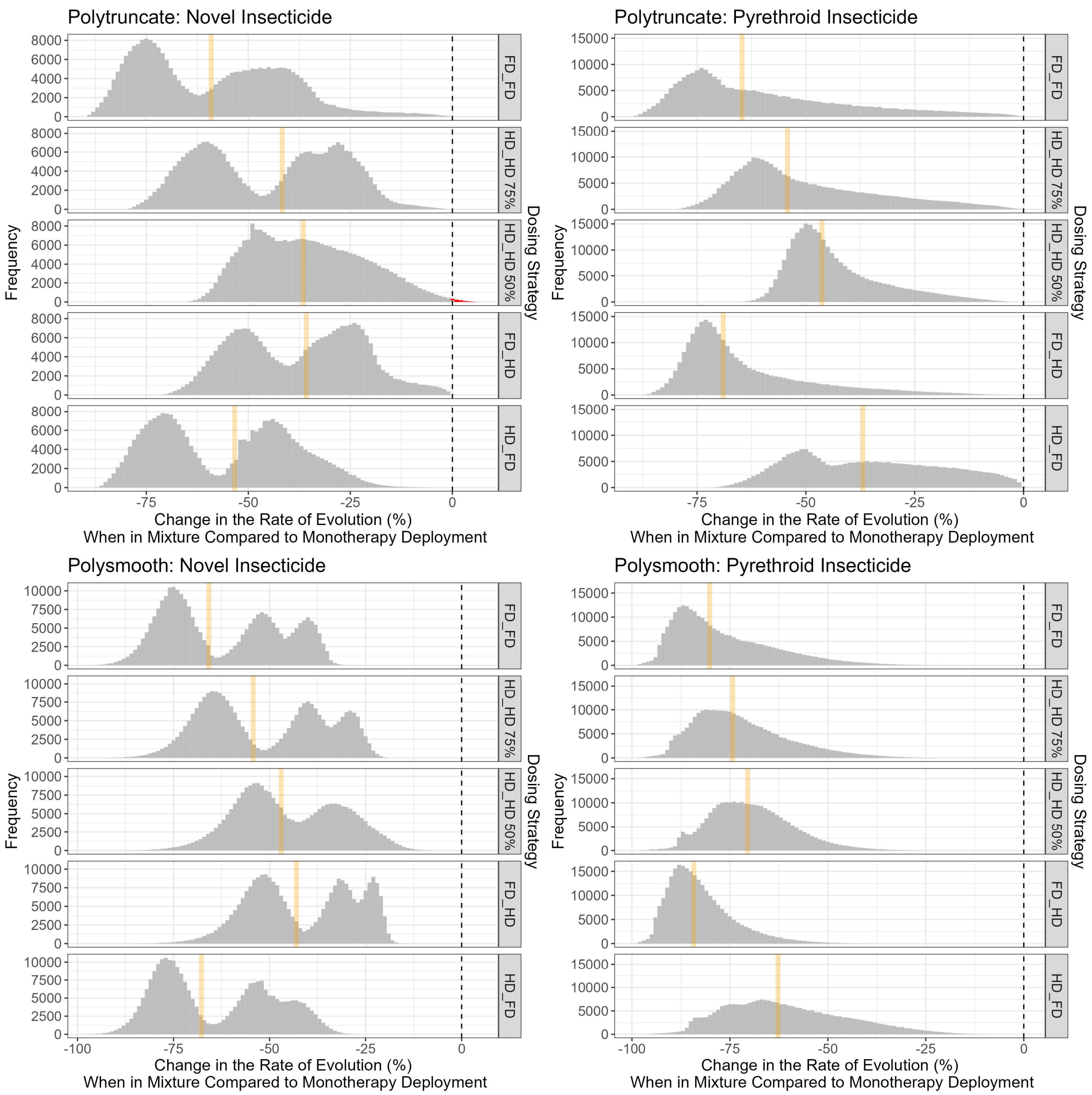


**Figure S1.1 Comparing the Mixture Dosing Strategies versus Monotherapy Deployments Stratified by Dosing Strategy.** Left panels is comparing each mixture dosing strategy versus deploying the novel insecticide monotherapy at full dose. The right panel is comparing each mixture dosing strategy versus deploying the pyrethroid insecticide in monotherapy. Green line indicate the mean value, orange line indicates the median value. Red bars indicate simulations where deploying as mixture was found to be the worse strategy. Dosing strategies are in the form Novel_Pyrethroid, with FD=full dose and HD = half dose. The further left values are, the greater the benefit of deploying the insecticides in the mixture versus monotherapy deployment.

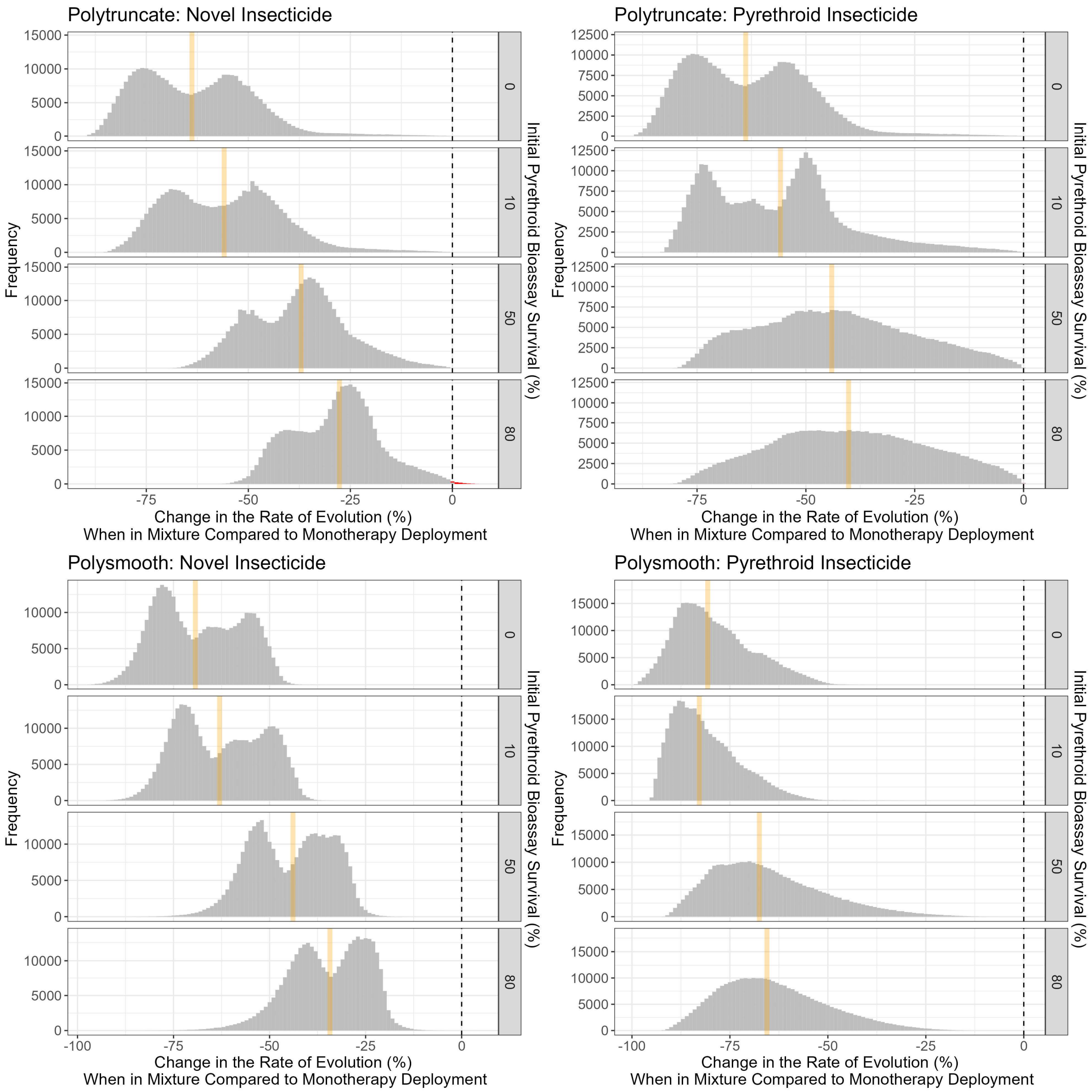


**Figure S1.2 Comparing mixture deployments versus monotherapy deployments stratified by initial bioassay survival to the pyrethroid.** Left Panel is comparing each mixture dosing strategy versus deploying the novel insecticide monotherapy at full dose. The right panel is comparing each mixture dosing strategy versus deploying the pyrethroid insecticide in monotherapy. Green line indicate the mean value, orange line indicates the median value. Red bars indicate simulations where deploying as mixture was found to be the worse strategy. Each panel indicates how much pyrethroid resistance there was at the start of the simulation, measured as bioassay survival (%).

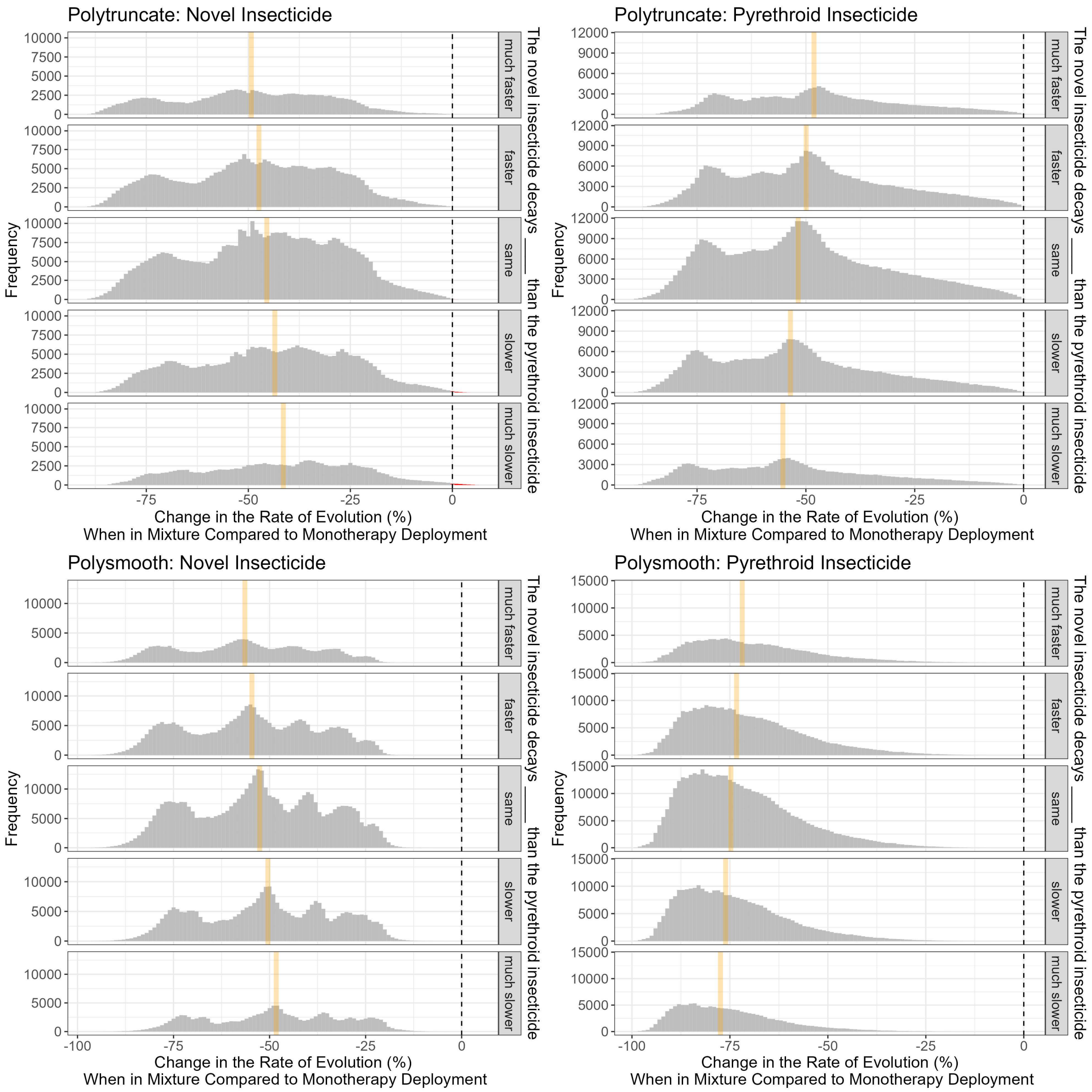


**Figure S1.3 Comparing mixture deployments versus monotherapy deployments stratified by decay rates.** Left Panel is comparing each mixture dosing strategy versus deploying the novel insecticide monotherapy at full dose. The right panel is comparing each mixture dosing strategy versus deploying the pyrethroid insecticide in monotherapy. Green line indicate the mean value, orange line indicates the median value. Red bars indicate simulations where deploying as mixture was found to be the worse strategy. The description at the side of each panel describes how the novel insecticide decays with respect to the pyrethroid, and therefore should be read as: The novel insecticide decays "description" than the pyrethroid insecticide.

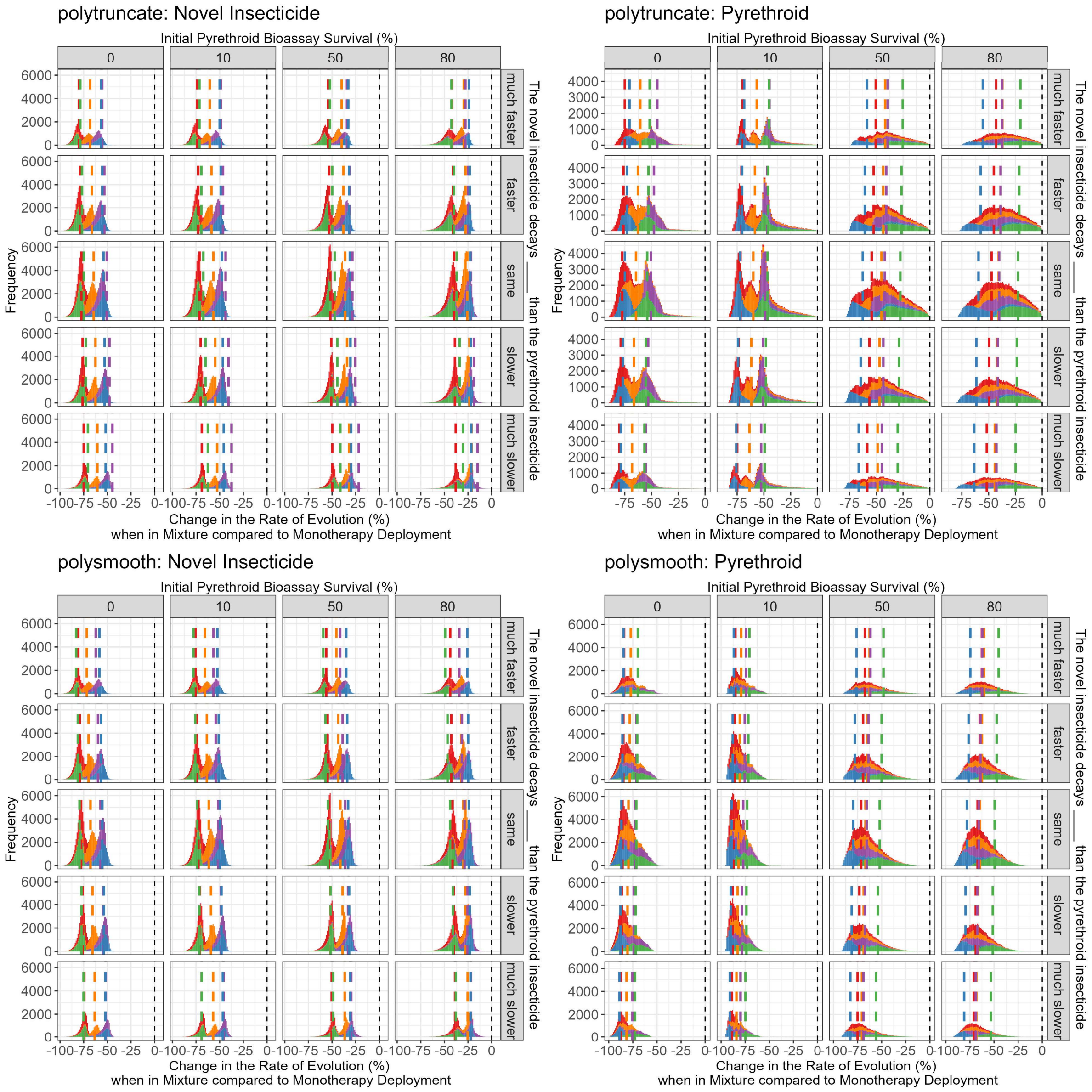


**Figure S1.4 Comparing mixture deployments versus so monotherapy lo deployments stratified by decay rates, initial pyrethroid resistance and mixture dosing.** Colours are the dosing strategies.

Red is full dose both insecticides. Purple is half dose of both insecticides. Green is full dose of the novel insecticide and a half dose of the pyrethroid.  Blue is the half dose of the novel insecticide and a full dose of the pyrethroid. Here, we are assuming a half dose retains 50% of the efficacy of a full dose. Stratification left to right is the resistance to the pyrethroid, and stratified top to bottom is the differences in decay rates. Top panels are the polytruncate model and bottom panels are the polysmooth panels. Left panels are the novel insecticide and the right panels are the pyrethroid insecticide.

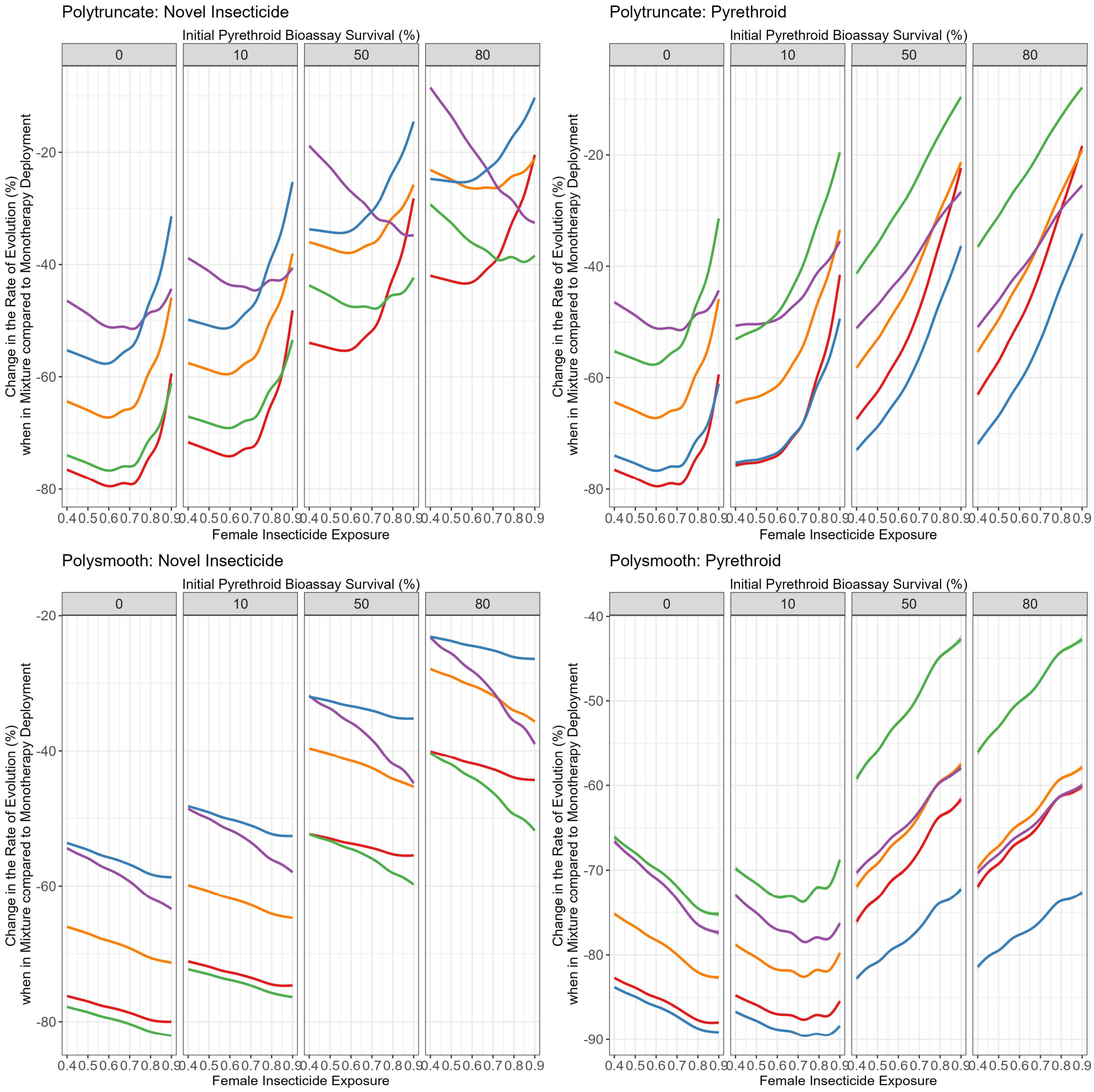


**Figure S1.5 Generalised Additive Model: Change in Rate of Evolution by Female Insecticide Encounter Probability.** Interpretation is that the lower the line, the better the dosing strategy is for slowing the rate of evolution. Left panels are for the novel insecticide, right panels are for the pyrethroid. Top panels look at polytruncate, and bottom panel looks at polysmooth. Additional generalised additive models smoothing over each parameter can be found in the supplementary information.

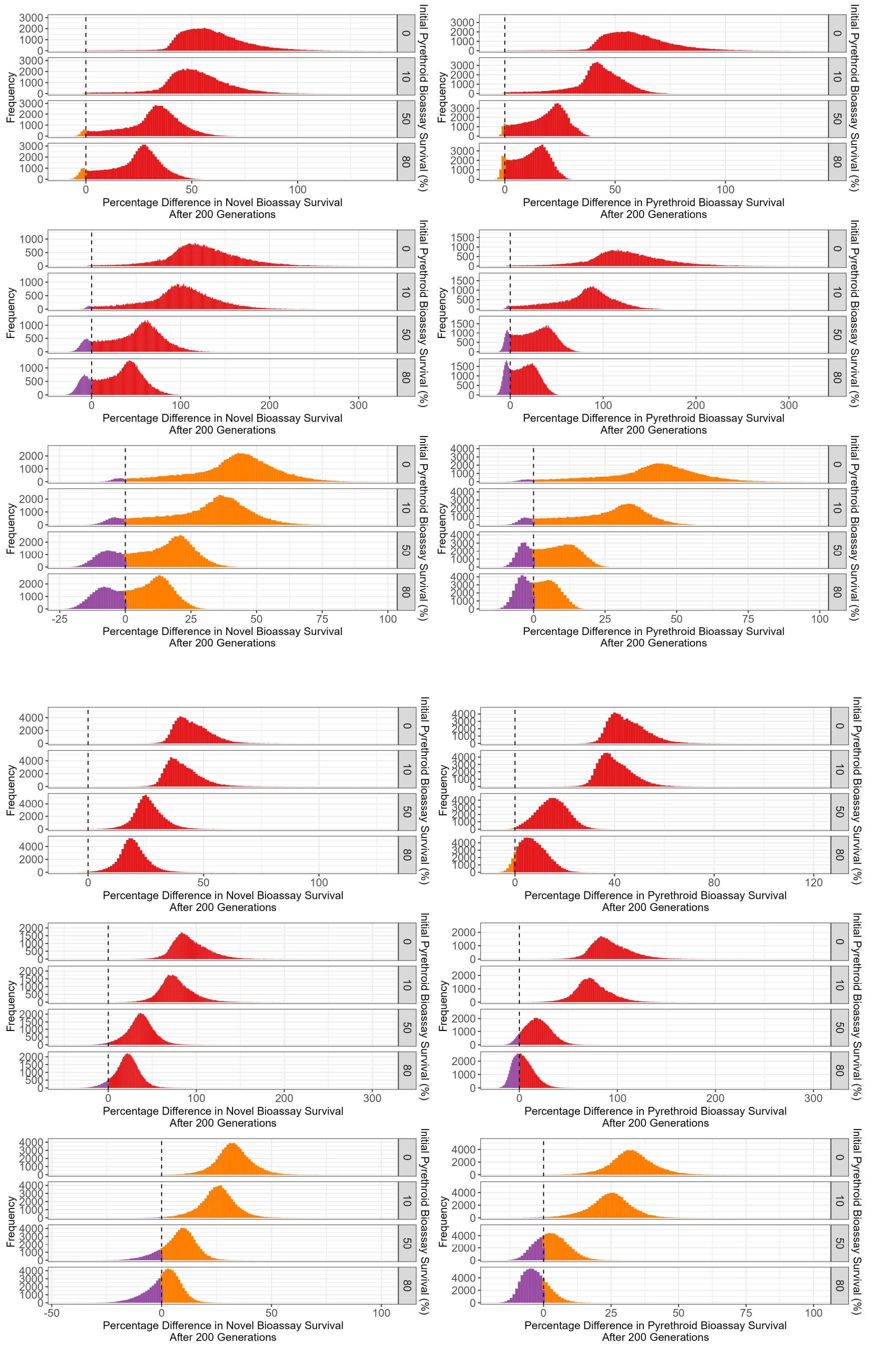


**Figure S1.6 Direct comparisons between mixture dosing strategies.** The colours indicate which dosing strategy performed better in the direct comparisons, where: Red = FD_FD, Purple = HD_HD retains 50% efficacy, Orange = HD_HD retains 75% efficacy. Top graphs (rows 1:3, in Green Border) are polytruncate model. Bottom 3 rows (4:6, in Blue Border) are polysmooth model.

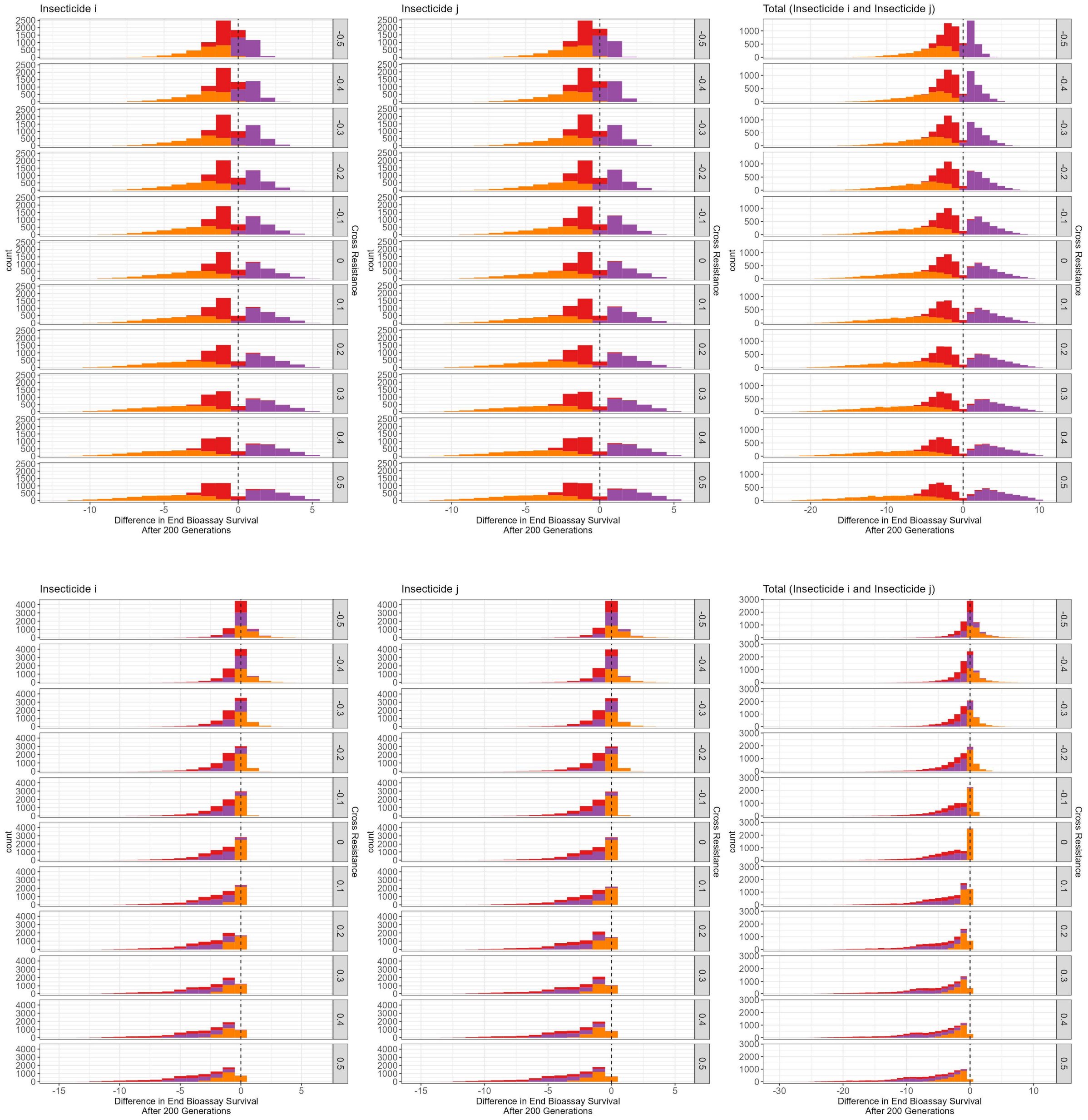


**Figure S1.7 Impact of cross resistance when deployed mixture versus deployed as a monotherapy sequence.** Top graph panel: Polytruncate, Bottom graph panel: Polysmooth. Left panel is the change between the monotherapy deployed insecticide (insecticide $i$) and the insecticide when in mixture. Middle panel is the change in the change between the not deployed insecticide (insecticide $j$) and the insecticide when deployed in mixture. Right panel is the total change in bioassay survival.

Red = FD_FD, purple = HD_HD retains 50%, orange = HD_HD retains 75%. Rows are the degree of cross resistance between the insecticides.

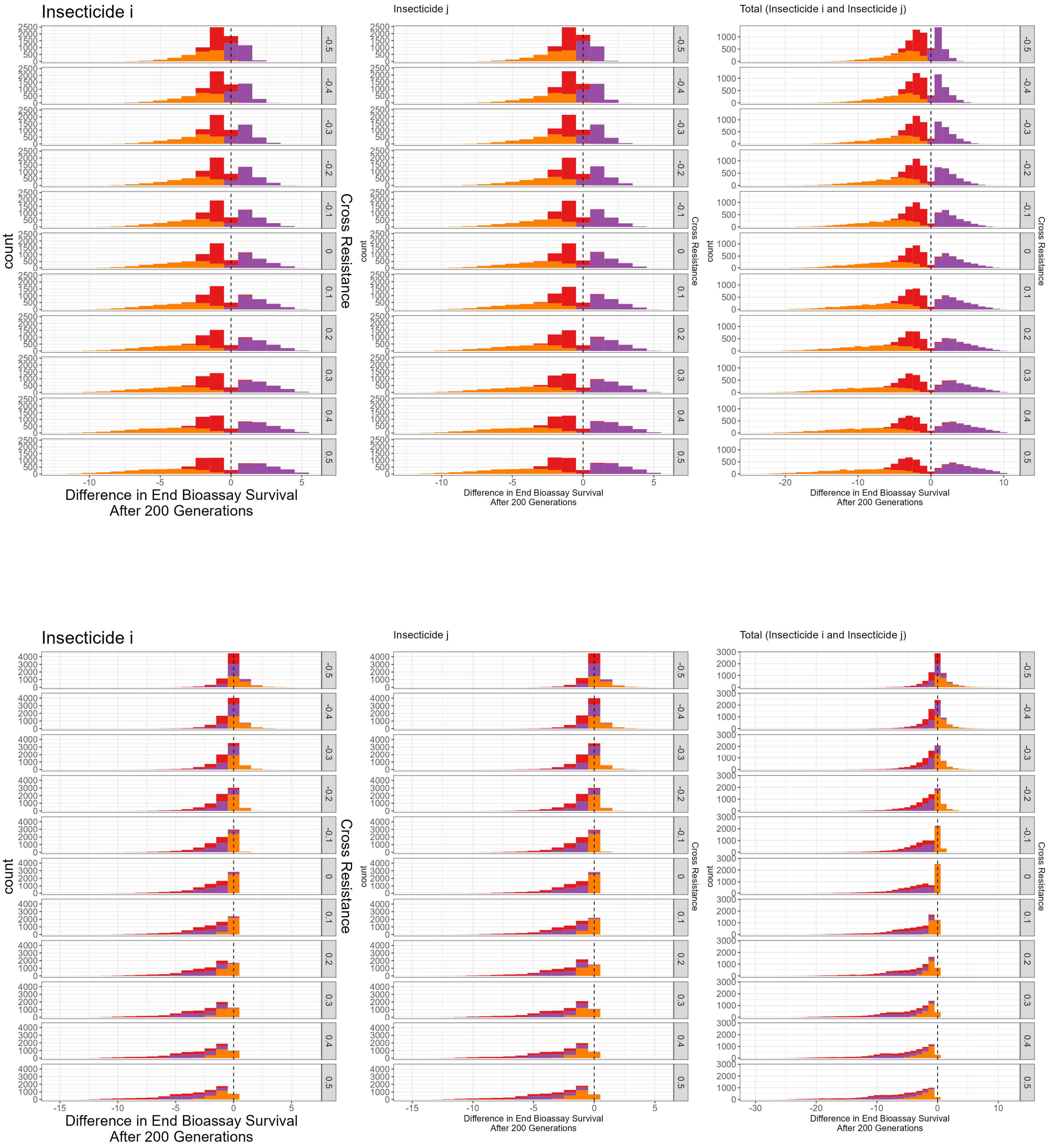


**Figure S1.8 Impact of Cross Resistance on the efficacy of mixtures versus monotherapy rotations to slow the rate of evolution of insecticide resistance**. Top graph panel: Polytruncate, Bottom graph panel: Polysmooth. Graphs are stratified by the amount of cross resistance between the two insecticides. Colours of the bars in indicate the dosing strategy of the mixture, where Red = both insecticides at full dose, orange = both insecticides at half dose and retain 75% of the full dose efficacy, and purple = both insecticides at half dose and retain 50% of their full dose efficacy.

| **Table S1: Comparing “polytruncate” and “polysmooth”.** | |
|---|---|
| Figure | Similarities/Differences |
| S1.1: Comparing the Mixture Dosing Strategies versus Monotherapy Deployments Stratified by Dosing Strategy | Main difference is there for “polytruncate” some simulations where deploying as a mixture is worse than deploying as a monotherapy, occurring only in the HD_HD (retains 50% ) simulations. Broad pattern is otherwise similar between both model branches. “Polytruncate” generally indicates a lower benefit of mixtures versus monotherapies than “polysmooth”. |
| S1.2 Comparing mixture deployments versus Monotherapy deployments stratified by initial bioassay survival to the pyrethroid. | Main difference is there for “polytruncate” some simulations where deploying as a mixture is worse than deploying as a monotherapy, occurring only in the initial bioassay survival to the pyrethroid is 80% simulations. Broad pattern is otherwise similar between both model branches. “Polytruncate” generally indicates a lower benefit of mixtures versus monotherapies than “polysmooth”. |
| S1.3 Comparing mixture deployments versus Monotherapy deployments stratified by decay rates. | Broad pattern is similar between both model branches. “Polytruncate” generally indicates a lower benefit of mixtures versus monotherapies than “polysmooth”. |
| Comparing mixture deployments versus Monotherapy deployments stratified by decay rates, initial pyrethroid resistance and mixture dosing. | Broad pattern is similar between both model branches. “Polytruncate” generally indicates a lower benefit of mixtures versus monotherapies than “polysmooth”. |
| S1.5 Generalised Additive Model: Change in Rate of Evolution by Female Insecticide Encounter Probability. | For “Polytruncate” increasing the female insecticide exposure decreases the decreases the benefit of the mixture versus the monotherapies. For “polysmooth” increasing the female insecticide exposure increases the benefit of the mixture versus pyrethroid monotherapies. And increasing the female insecticide exposure decreases the benefit of the mixture versus novel monotherapies when there is high resistance to the pyrethroid.This difference can be explained by the mechanism of selection. Where for “polytruncate” at high female insecticide exposure only very resistance individuals survive. Whereas for polysmooth because survival is dependent on the individual polygenic resistance score, even less resistant individuals have a chance to survive. |
| S1.6 Direct comparisons between mixture dosing strategies | Broad pattern is similar between model branches. |
| S1.7 Impact of cross resistance when deployed mixture versus deployed as a Monotherapy sequence. | Difference is that when there is positive cross resistance and a HD_HD (retains 50%) this is worse for the mixture in the “polytruncate” simulations while marginally better in the “polysmooth” simulations.<br>The broad pattern of FD_FD > HD_HD (retains 75%) > HD_HD (retains 50%) is consistent between both model branches. |
| S1.8 Impact of Cross Resistance on the Efficacy of Mixtures versus Monotherapy Rotations to slow the rate of evolution of insecticide resistance. | The models differ in that when there is positive cross resistance and a HD_HD (retains 50%) is compared against the monotherapy rotations, for “polytruncate” the mixture performs worse, whereas for “polysmooth” the mixture performs marginally better. The broad pattern of FD_FD > HD_HD (retains 75%) > HD_HD (retains 50%) is consistent between both model branches. |
