## Supplement 2 for "Exploring operational requirements of mixtures for insecticide resistance management in public health using a mathematical model assuming polygenic resistance"

### Supplement 2: Comparison between “polytruncate” and “polysmooth”

While the polytruncate and polysmooth models clearly differ from one another (e.g., Figures S1.5 and S1.8; Supplement 1), it is also important to understand how (and if) the broad conclusions from the model are impacted. The simplest way to look at this is to see if the FD_FD mixture or HD_HD (retaining 75% or retaining 50% efficacy) mixture performed better (Figure 7) but then directly compare if the outcomes from both models give the same conclusion, and if not what parameter(s) appear to drive this difference; and whether this can be explained by the change in the assumption of how selection is assumed to occur in the model. Figures S2.1-S2.3 shows that the models diverge (in terms of dosing strategy recommendation) at higher levels of resistance. We see that agreement between the models is very high when resistance to both insecticides is low. When resistance to the pyrethroid is high, there is lower agreement between the models. Agreement between the models was generally lowest when comparing the 75% and 50% mixtures against one another. Model agreement was best when comparing the full-dose mixture versus the 75% mixture.

When looking at cross resistance and dosing, it can be seen the models diverge in whether the mixture strategy or monotherapies is better (Figure S2.5, top row). However, it can be seen the magnitude of these differences is frequently small (Figure S2.5, bottom row).

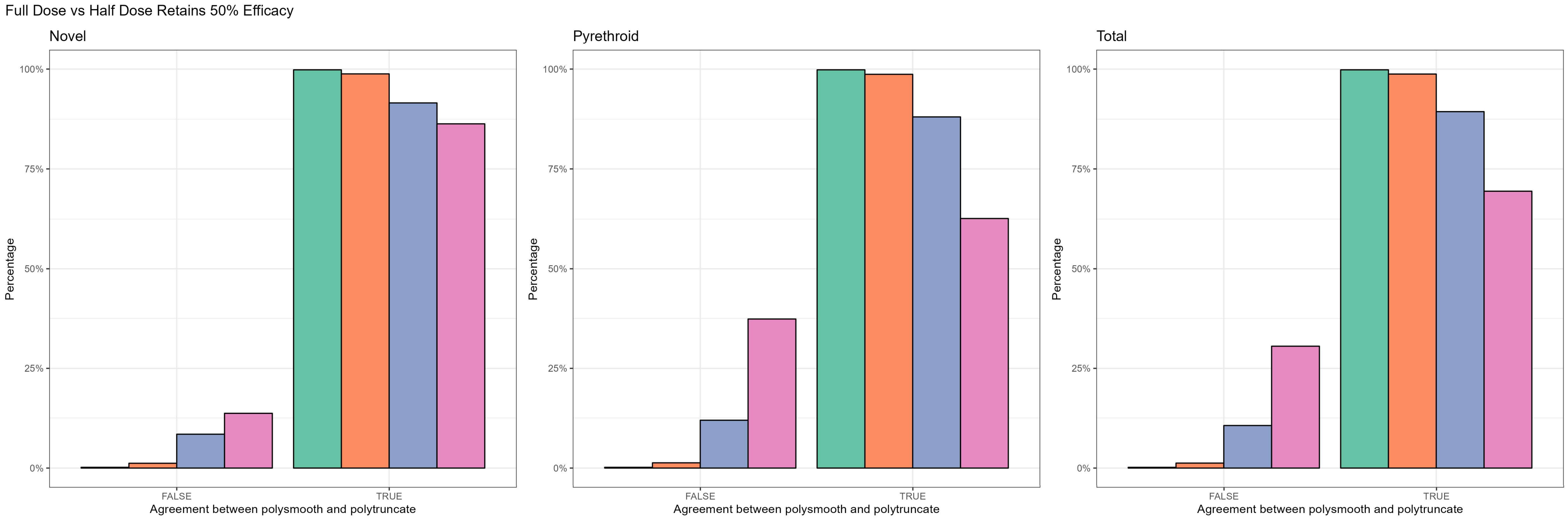


**Figure S2.1: Comparing the Outcome Between polytruncate and polysmooth for Full-Dose Mixtures vs Half Dose Mixtures Retaining 50% Efficacy.** False = The models did not agree which dosing strategy was best. True = The models did agree which dosing strategy was best. Colours indicate the initial amount of resistance to the pyrethroid measured in a bioassay: teal = 0%, salmon = 10%, blue = 50%, pink = 80%. Each plot indicates whether there was an agreement for just the novel (left), just the pyrethroid (centre) or both (right).

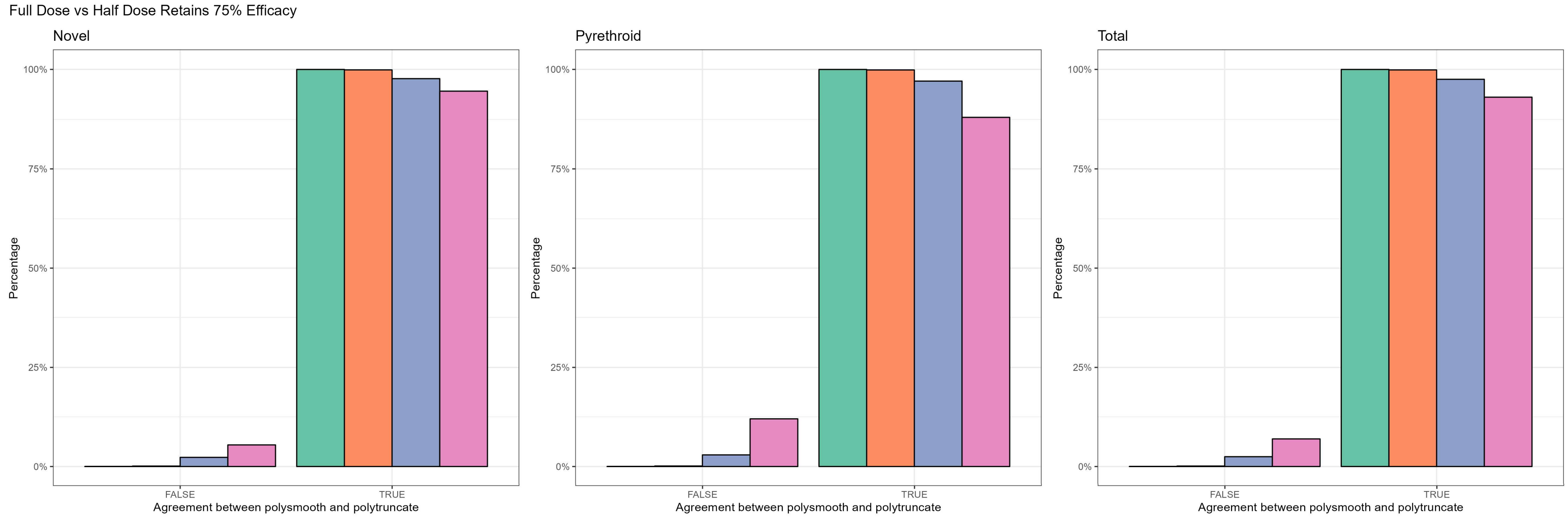


**Figure S2.2: Comparing the Outcome Between polytruncate and polysmooth for Full-Dose Mixtures vs Half Dose Mixtures Retaining 75% Efficacy.** False = The models did not agree which dosing strategy was best. True = The models did agree which dosing strategy was best. Colours indicate the initial amount of resistance to the pyrethroid measured in a bioassay: teal = 0%, salmon = 10%, blue = 50%, pink = 80%. Each plot indicates whether there was an agreement for just the novel (left), just the pyrethroid (centre) or both (right).

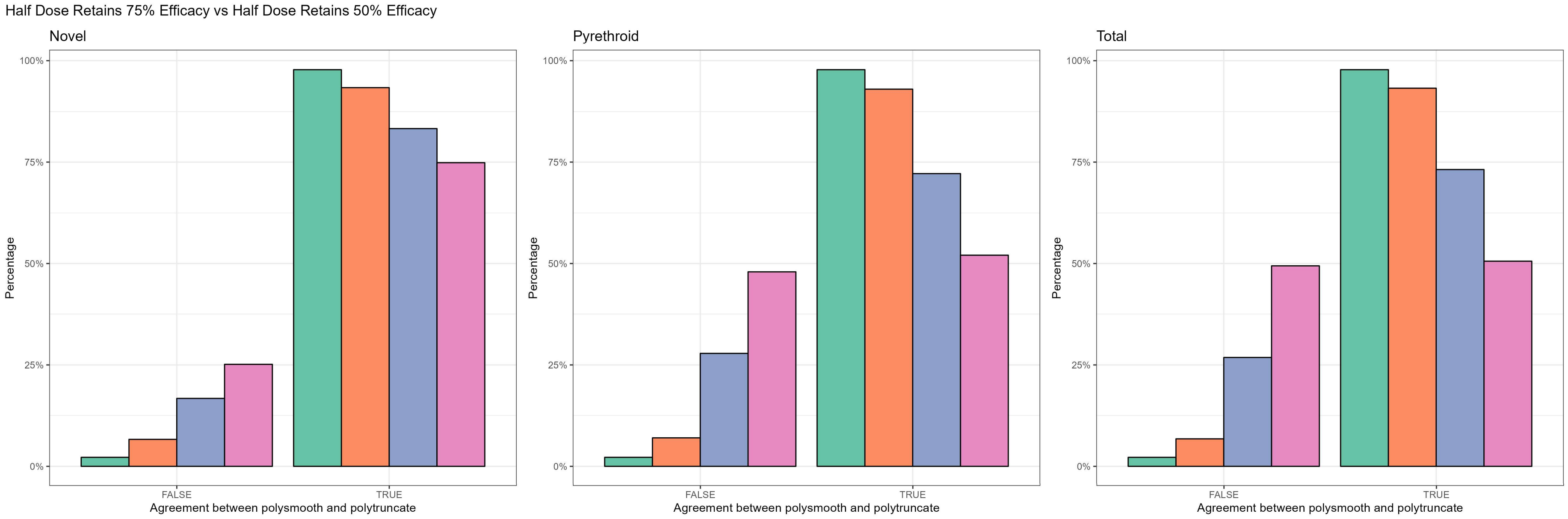


**Figure S2.3 Comparing the Outcome Between polytruncate and polysmooth for Haf-Dose Mixtures retaining 75% Effacacy vs Half Dose Mixtures Retaining 50% Efficacy.** False = The models did not agree which dosing strategy was best. True = The models did agree which dosing strategy was best. Colours indicate the initial amount of resistance to the pyrethroid measured in a bioassay: teal = 0%, salmon = 10%, blue = 50%, pink = 80%. Each plot indicates whether there was an agreement for just the novel (left), just the pyrethroid (centre) or both (right).

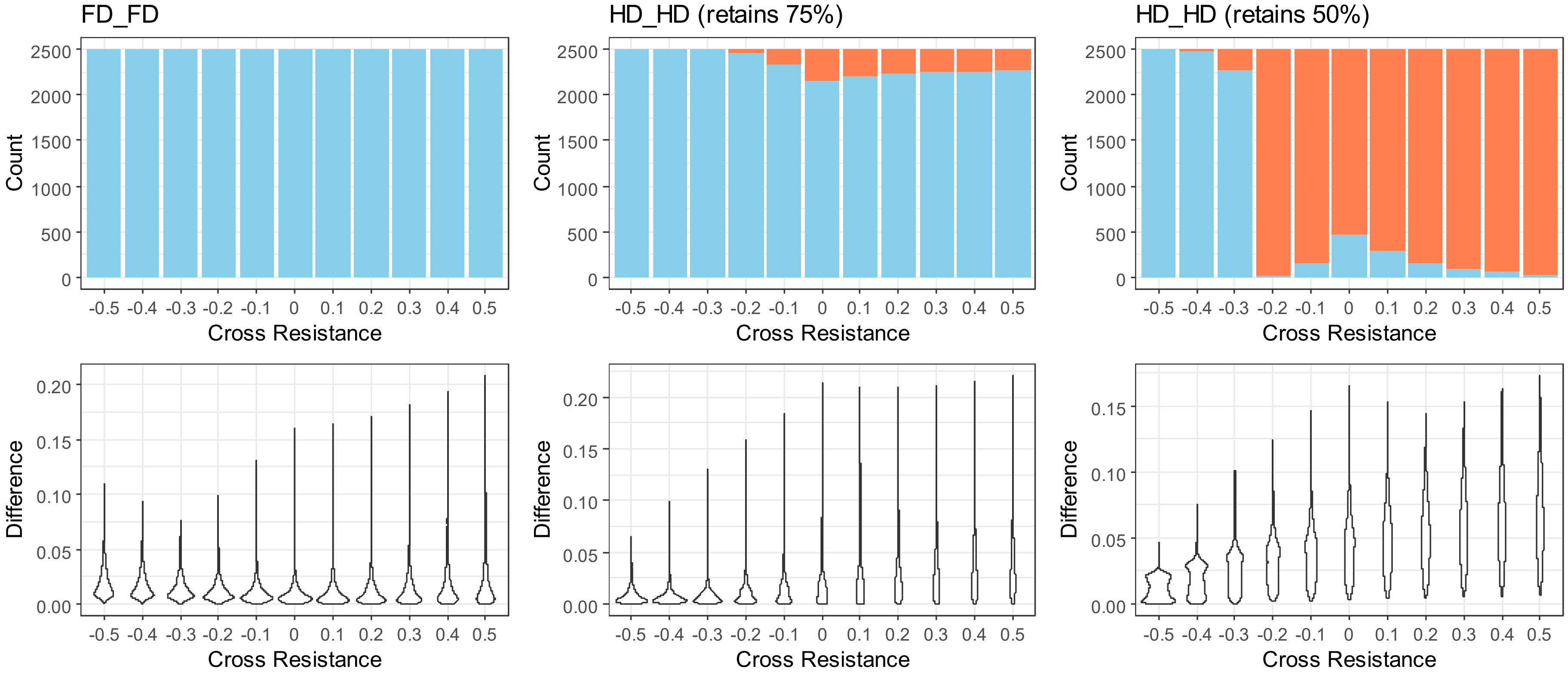


**Figure S2.4: Model Agreement when comparing mixtures versus a single monotherapy sequence.** Top row: Model agreement as to whether the mixture strategy was better or not (blue = mixtures better, red = mixtures worse). The bottom row is the magnitude of these differences (as violin plots).

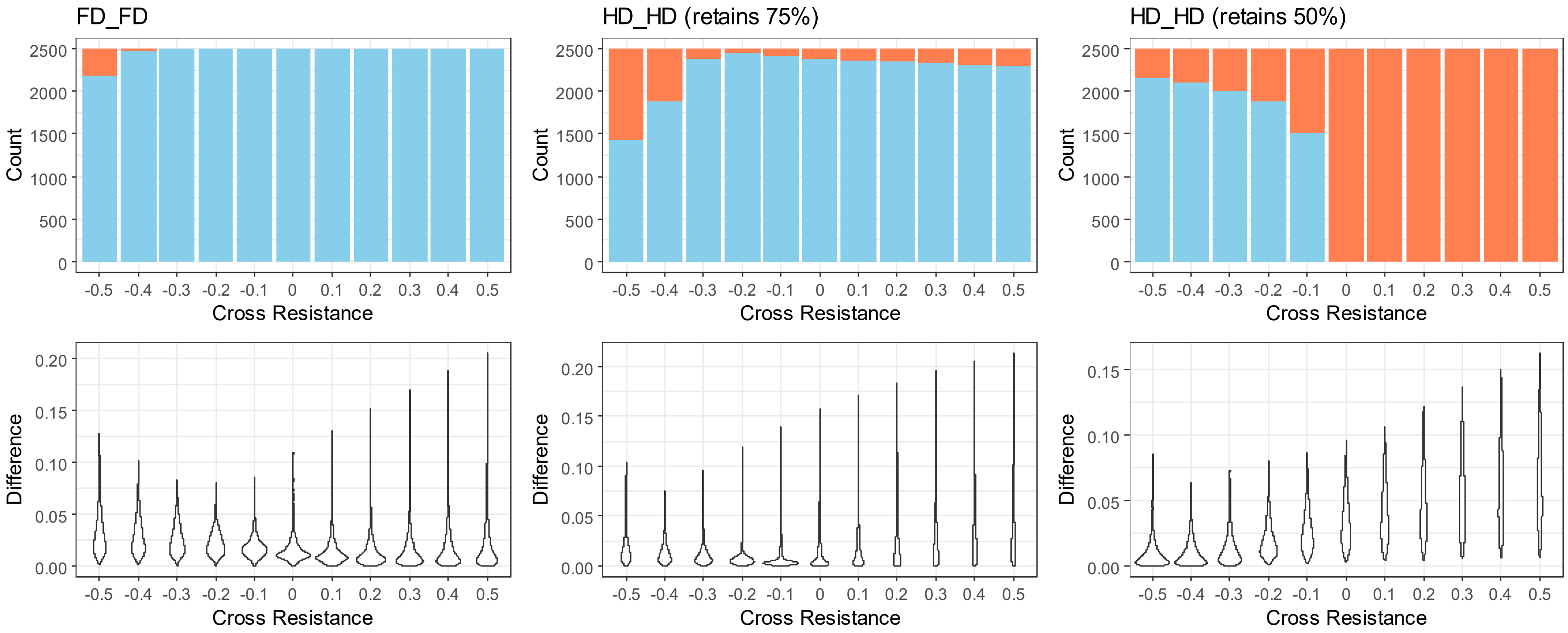


**S2.5 Model Agreement when comparing mixtures versus monotherapy rotations.** Top row: Model agreement as to whether the mixture strategy was better or not (blue = mixtures better, red = mixtures worse). The bottom row is the magnitude of these differences.
